# Decoding super-enhancer regulatory grammar and pathogenic gain-of-function variants directly from DNA sequence

**DOI:** 10.64898/2026.10.11.756618

**Authors:** Yan Liang, Xiaohui Xu, Yiping Mao, Zehua Zheng, Nan Liu

**Affiliations:** Liangzhu Laboratory, Zhejiang University School of Medicine, Hangzhou, Zhejiang, China; Institute of Hematology, Zhejiang University, Hangzhou, Zhejiang, China; Bone Marrow Transplantation Center of the First Affiliated Hospital, Zhejiang University School of Medicine, Hangzhou, Zhejiang, China

## Abstract

Super-enhancers (SEs) drive cell-identity gene expression, yet their identification relies on epigenomic profiling (e.g., H3K27ac ChIP-seq), precluding discovery in unprofiled cellular states and leaving their underlying sequence grammar unknown. Here, we present Borzoi-SE, a deep learning framework built upon the genomic transformer Borzoi that formulates SE discovery as hierarchical, cell-type-resolved genomic object detection directly from long-range sequence across 90 human lineages. Base-resolution attribution recovers lineage-defining master transcription factors across cell types. Within SEs shared across lineages, it reveals lineage-dependent motif usage, suggesting that the same regulatory regions can support distinct cell-type-specific programs. Using *in silico* ablation, Borzoi-SE decoded the functional hierarchy of the β-globin super-enhancer, prioritizing hypersensitive site 2 (HS2) as the dominant constituent, which we experimentally verified via CRISPR interference. Trained without variant-effect labels, Borzoi-SE identified gain-of-super-enhancer variants in different tissues. At the *CHRM2* locus, a single-nucleotide variant created a *de novo* glucocorticoid receptor (GR) motif that recruited activated GR and potentiated adjacent MEF2 regulatory syntax, nucleating a cardiac super-enhancer. Orthogonal multi-omics and population-genetic evidence linked this variant to increased *CHRM2* expression and variation in resting heart rate and PR-interval duration. Borzoi-SE establishes a general framework for decoding SE regulatory grammar and prioritizing causal non-coding variants directly from sequence.

## Introduction

Super-enhancers (SEs) are large, densely clustered *cis*-regulatory domains that direct dense assemblies of transcription factors (TFs) to control the expression of genes governing cell identity and disease^1^. Despite their fundamental biological importance, the identification of SEs has remained tethered to empirical epigenomic profiling: the canonical ROSE algorithm stitches experimental H3K27ac or MED1 ChIP-seq peaks across arbitrary distance thresholds, retaining only those domains that surpass an empirical signal cutoff^1^. This experimental dependency restricts SE discovery strictly to profiled cell states, rendering the mapping of transient or rare cellular contexts prohibitive. More fundamentally, because peak-stitching algorithms are descriptive rather than generative, they merely catalogue pre-existing chromatin states without uncovering the underlying sequence grammar or master transcription factors that drive cluster formation. Consequently, they cannot decipher whether super-enhancer logic is encoded directly within primary DNA sequence, nor can they predict *de novo* super-enhancer nucleation or evaluate the functional impact of non-coding genetic variation.

While omics-based methods, such as ChIP-seq peak stitching and 3D chromatin conformation approaches, have successfully mapped active SEs in well-characterized systems, they remain constrained by high profiling costs, empirical distance thresholds, and an inability to interrogate sequence determinants directly^1–5^. To overcome these experimental dependencies, first-generation sequence-based models including DeepSE, SENet, MuSE, and DcSE have emerged^6–9^. However, their architectures did not reflect the multi-scale biology of SEs. First, these models frame SE discovery as binary classification of isolated sequence snippets (≤ 3 kb), typically processed as discrete *k*-mer composition vectors or shallow embeddings. Because authentic SEs are higher-order genomic domains spanning tens to hundreds of kilobases that arise from the spatial clustering of constituent enhancers and insulator boundaries, classifying truncated, out-of-context fragments fundamentally blinds these models to the long-range spatial syntax that separates true super-enhancers from isolated typical enhancers^1,10^. Second, existing models are cell-type agnostic: by pooling annotations across disparate tissues into a single binary scalar, they cannot account for the fact that SE activity is dynamically executed by lineage-specific master TFs, or that shared regulatory domains often rely on distinct regulatory grammars across tissues. Third, these classifiers have relied almost universally on random genomic partitioning, allowing homologous regulatory sequences to bleed between training and test sets and confounding genuine sequence learning with genomic memorization^7^. Existing tools can at best ask whether a short fragment is “enhancer-like”, but remain incapable of pinpointing where an SE begins and ends, resolving its cell-type-specific activity, or identifying the cooperative cis-regulatory elements that establish domain potency.

In parallel, the emergence of sequence-to-function foundation models, including Basenji, Enformer, PromoterAI, Borzoi and AlphaGenome, has transformed regulatory genomics by predicting continuous, base-resolution epigenomic and transcriptomic profiles (e.g., CAGE, ChIP-seq, and RNA-seq) directly from DNA sequence across hundreds of cell types^11–15^. While remarkably adept at functional track regression, these architectures do not directly demarcate physical boundaries or model the cooperative clustering that characterizes super-enhancers. Furthermore, coupling their predicted H3K27ac profiles with empirical peak-stitching tools (e.g., ROSE) inevitably compounds prediction noise and heuristic thresholds across chromosome-scale inputs. Harnessing the rich latent representations learned by sequence transformers to directly predict and structurally resolve higher-order regulatory domains therefore represents a critical yet unrealized transition from continuous track regression to functional entity discovery.

Here, we introduce Borzoi-SE, a deep learning framework that formulates super-enhancer discovery directly from primary sequence as hierarchical, cell-type-resolved genomic object detection. Built upon the Borzoi foundation model, which internalizes regulatory priors from thousands of functional genomics assays, Borzoi-SE operates over 524-kb genomic inputs to capture the broad chromatin topology and distant boundaries characteristic of super-enhancers^13^. Rather than treating SE discovery as a cell-type-agnostic classification of isolated fragments, Borzoi-SE integrates cell-type-resolved prediction with end-to-end genomic object detection. This is achieved through two specialized modules: a dilated Cluster head (∼64.7 kb) that models long-range constituent enhancer clustering to demarcate physical domain boundaries and suppress background noise, and an SE head that evaluates cooperative motif syntax to simultaneously resolve cell-type-specific super-enhancer activity across 90 human cell and tissue lineages.

To evaluate model generalizability without chromosome-specific bias, we established a five-fold cross-chromosome partitioning scheme (independently training models across distinct chromosome combinations) and benchmarked Borzoi-SE against existing fragment-based classifiers. Borzoi-SE not only matches baseline discriminative fidelity on isolated fragments but substantially outperforms them in genome-wide boundary delineation and false-positive suppression, while accurately ranking continuous regulatory potency across all lineages. Beyond locus-level detection, base-resolution gradient attribution enables *de novo* recovery of master transcription factor syntax and reveals that shared regulatory domains dynamically switch functional motif usage across cell states.

Crucially, a unique strength of sequence-based deep learning is its capability to perform high-throughput *in silico* saturation mutagenesis, systematically simulating genetic perturbations and evaluating higher-order cooperativity directly on primary sequence. At the human *β*-globin locus, *in silico* constituent ablation identified HS2 as the preeminent master hub whose disruption triggers a domain-wide collapse of chromatin accessibility and downstream expression, a functional hierarchy we experimentally validated in living cells via CRISPR interference (CRISPRi). Furthermore, Borzoi-SE accurately models pathogenic gain-of-super-enhancer mutations ab initio, including the classic somatic insertion activating *TAL1* in T-cell acute lymphoblastic leukemia (T-ALL) and related hematopoietic lineages, and successfully discovers previously uncharacterized non-coding gain-of-SE variants that nucleate *de novo* cardiac enhancers via the creation of transcription factor motifs^16^. By prioritizing causal variants calibrated against population-scale histone acetylation QTLs (haQTLs) and orthogonal multi-omics cardiac assays, Borzoi-SE establishes that cell-type-resolved, quantitative super-enhancer landscapes can be mapped, interpreted, and perturbatively interrogated from sequence alone without experimental chromatin profiling.

## Results

### Borzoi-SE enables cell-type-specific super-enhancer discovery through hierarchical genomic object detection

To overcome the reliance of conventional methods (such as ROSE) on experimental peak-stitching, we formulated super-enhancer (SE) discovery as a sequence-based genomic object detection task and developed Borzoi-SE, a deep learning framework that directly maps long-range DNA sequence to cell-type-specific SE domains (**Fig. 1a**). Given a 524-kb DNA sequence, Borzoi-SE extracts sequence representations via a pretrained Borzoi foundation backbone down-sampled to a 256-bp spatial resolution across a central ∼410-kb window (**Fig. 1a**, Module 1; Methods). To mirror the biological logic of enhancer stitching, Borzoi-SE routes these representations through two parallel branches: one preserving local sequence context and the other driving a dedicated Cluster head with a 64.7-kb receptive field (253 bins), which accommodates the classical 25-kb stitching window of ROSE to model cluster nucleation while suppressing background noise (**Fig. 1a**, Module 2; Methods). A Squeeze-and-Excitation channel attention module then adaptively recalibrates context features against cluster logits, allowing an SE head to evaluate localized transcription factor (TF) motif syntax against broader cluster topology (**Fig. 1a**, Modules 3 and 4; Methods). Unlike traditional binary sequence classifiers, Borzoi-SE outputs a cell-type-by-position prediction matrix (90 cell-type channels × 1,600 bins across the central ∼409.6 kb at 256-bp resolution), simultaneously predicting the quantitative SE probability and physical coordinates across 90 human cell and tissue lineages (**Fig. 1a, b, Supplementary Table 1**).

**Figure 1:**
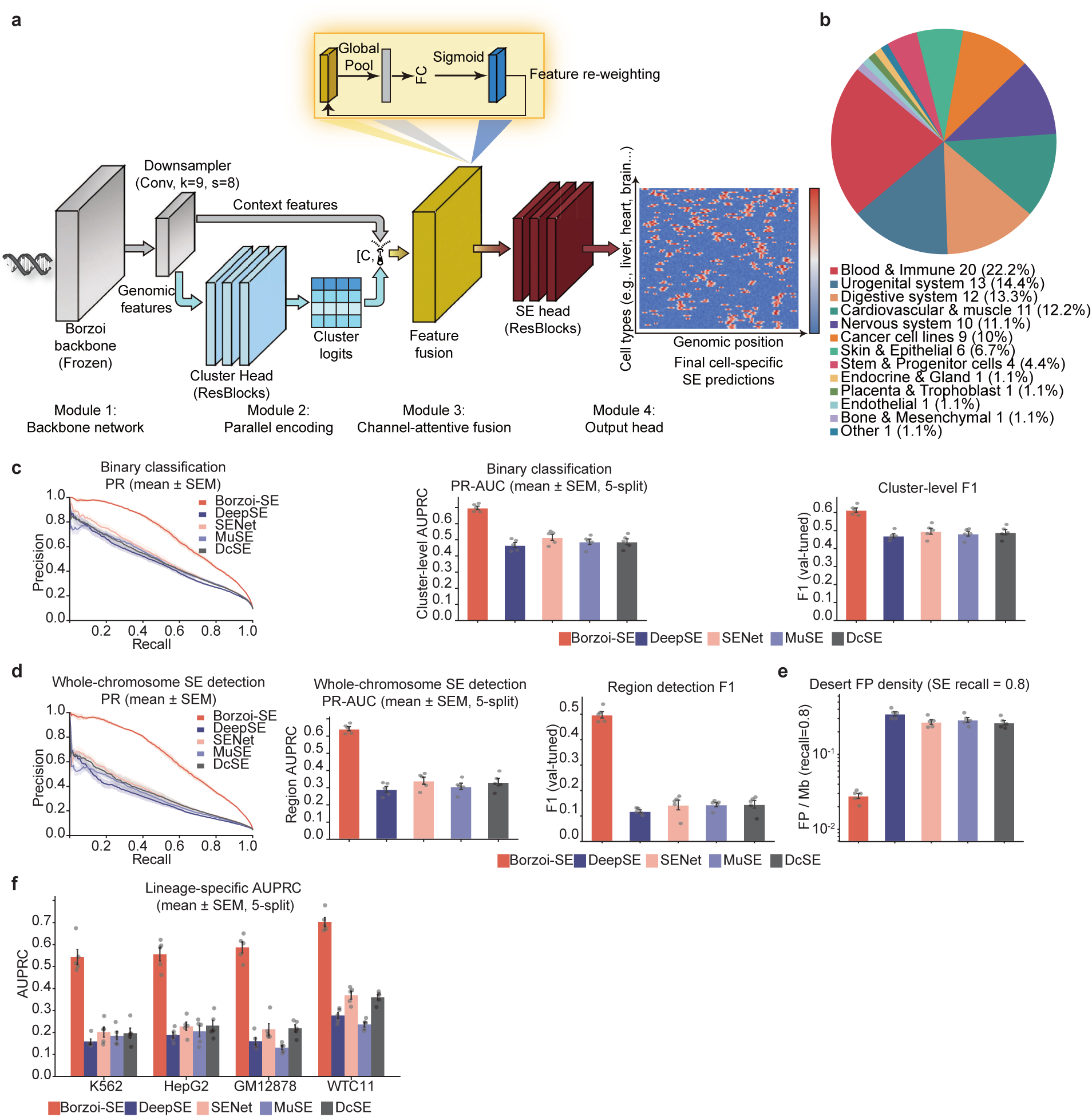
Overview of the Borzoi-SE architecture and cross-chromosome benchmark evaluation. **a**, Schematic of the Borzoi-SE framework, highlighting the Borzoi backbone (Module 1), dilated Cluster Head (Module 2), channel-attentive feature fusion (Module 3), and SE head (Module 4). **b**, Composition of the 90 modeled human cell and tissue types grouped by physiological system. **c**, Pan-tissue binary classification on pre-stitched enhancer clusters (Experiment 1): precision–recall (PR) curves (left), AUPRC (middle), and validation-tuned F1 scores (right). **d**, Whole-chromosome *de novo* SE detection on held-out test chromosomes (Experiment 2): region-level PR curves (left), AUPRC (middle), and region F1 scores (right; IoU ≥ 0.3). **e**, False-positive density across quiescent genomic deserts (FP/Mb, log scale) at 80% SE recall. **f**, Lineage-specific AUPRC across four representative cell lines comparing Borzoi-SE with retrained baseline architectures (Experiment 3). For **c**–**f**, solid lines and bar heights denote the mean across *n* = 5 independent chromosome splits; shaded ribbons and error bars denote s.e.m.; individual points represent separate splits.

We benchmarked Borzoi-SE against four published models using five independent chromosome splits, each with two held-out test chromosomes (Splits 0–4; Methods)^6–9^. All models were independently trained, validated, and tested across all five splits to ensure robust genome-wide generalization. We first evaluated pan-tissue binary classification on complete, variable-length ROSE-stitched enhancer clusters (Experiment 1; Methods). Test sets preserved the natural genomic ratio of SEs to typical enhancer clusters (TEs; ∼1:10) without artificial down sampling. Across all five chromosome splits, Borzoi-SE substantially outperformed all four baseline architectures. It elevated the PR-AUC from baseline plateaus of 0.468–0.516 to 0.698 and increased the validation-tuned F1 score from 0.471–0.497 to 0.614 (two-sided paired t-test, all p<0.01; **Fig. 1c, Supplementary Table 2**). Borzoi-SE also maintained consistently superior discrimination in ROC-AUC (0.938 vs. 0.883–0.895; **Supplementary Fig. 1a**). These results indicate that even in pan-tissue binary classification of pre-segmented enhancer clusters - the setting for which DeepSE, SENet, MuSE, and DcSE were designed - Borzoi-SE substantially outperformed these specialized models, highlighting the advantage of long-range contextual modeling^6–9^.

While discrete segment classification assesses sequence discrimination, authentic super-enhancers are continuous domains spanning dozens to hundreds of kilobases. In realistic discovery settings, regulatory boundaries are not known *a priori*. Because existing baseline models are restricted to pre-cropped 3-kb fragments, they cannot determine where an SE begins or ends, nor can they prevent false alarms across vast non-coding expanses. To evaluate *de novo* discovery in an unconstrained setting, we challenged all models with pure end-to-end genomic object detection across continuous, unfragmented held-out test chromosomes (Experiment 2; Methods). Following whole-chromosome sliding scans with assembly gaps and ENCODE blacklists masked, predicted regions were stitched (≤ 12.5 kb) and matched to ground-truth Union SEs using greedy one-to-one matching (IoU ≥ 0.3)^17^. Borzoi-SE doubled the region-level detection precision of baseline models, achieving a PR-AUC of 0.643 and an F1 score of 0.498, compared to baseline ceilings of ∼0.34 PR-AUC and ∼0.15 F1 (two-sided paired *t*-test, all *p* < 10^−4^; **Fig. 1d, Supplementary Table 2**). Across extensive quiescent genomic deserts (unannotated background regions outside known enhancer clusters), where localized models generated persistent false-positive noise (0.265–0.349 FP/Mb), Borzoi-SE suppressed false alarms by approximately tenfold (0.028 ± 0.002 FP/Mb at 80% recall; **Fig. 1e**). Moreover, among true-positive detections at this recall, Borzoi-SE demarcated physical boundaries with high spatial precision (median IoU of 0.668 versus 0.479–0.502 for baseline models; **Supplementary Fig. 1b**). These findings demonstrate that distal sequence context is indispensable for suppressing spurious background activations and accurately resolving domain boundaries on continuous chromosomes.

Finally, we asked whether models could resolve cell-type-specific regulatory grammar across four representative cellular contexts spanning distinct developmental lineages: K562 (erythroid), HepG2 (endodermal epithelium), GM12878 (lymphoblastoid), and WTC11 (pluripotent stem cells; experiment 3; Methods). To ensure a fair and direct comparison, all four baseline architectures were independently retrained for each individual lineage across all five splits (80 retrained models in total; Methods). The lineage-dedicated output channels of Borzoi-SE consistently outperformed these specialized baselines, attaining on-target PR-AUCs of 0.544–0.703 across all four cell types compared to 0.131–0.369 for the baselines (two-sided paired t-test, all p<0.01; **Supplementary Table 2**, **Fig. 1f**). While retrained baselines achieved moderate ROC-AUC scores (0.682–0.910; **Supplementary Fig. 1c**), their precision collapsed under natural genomic prevalence. This divergence confirms that short-range baseline models can distinguish active enhancer regions from quiescent genomic background (yielding moderate ROC-AUCs), but lack the long-range spatial context required to separate bona fide SEs from typical enhancers under natural genomic prevalence (causing precision collapse).

Although baseline models were independently retrained using lineage-specific labels, we asked whether they truly internalized lineage-defining regulatory grammar rather than shared enhancer features. We therefore evaluated cross-lineage prediction matrices across all cell-type pairs (4 × 4 heatmaps; **Supplementary Fig. 1d**). Borzoi-SE exhibited prominent diagonal dominance, with on-target PR-AUCs of 0.544–0.703 markedly exceeding off-target scores of 0.159–0.328, demonstrating strict lineage exclusivity. In stark contrast, baseline models produced indiscriminate cross-lineage profiles lacking diagonal enrichment (PR-AUC 0.114–0.369 across all combinations; **Supplementary Fig. 1d**). This discrepancy reveals that localized classifiers rely on shortcut learning from ubiquitous regulatory marks, whereas the hierarchical architecture of Borzoi-SE successfully captures the cooperative syntax that defines lineage-specific super-enhancer programs.

### Borzoi-SE delineates SE boundaries and quantitatively decodes continuous regulatory potency across human lineages

Evaluations on held-out test chromosomes demonstrated robust genome-wide performance, with a pooled region-level PR-AUC of 0.506 and an ROC-AUC of 0.895 across all 90 human cell types (**Fig. 2a, b**). Specifically, Borzoi-SE achieved cell-type-specific region-level PR-AUC scores ranging from **0.28 to 0.73 (median: 0.535)** and ROC-AUC scores from **0.75 to 0.95 (median: 0.902)** across 90 individual tissues and cell types (**Fig. 2c**).

**Figure 2:**
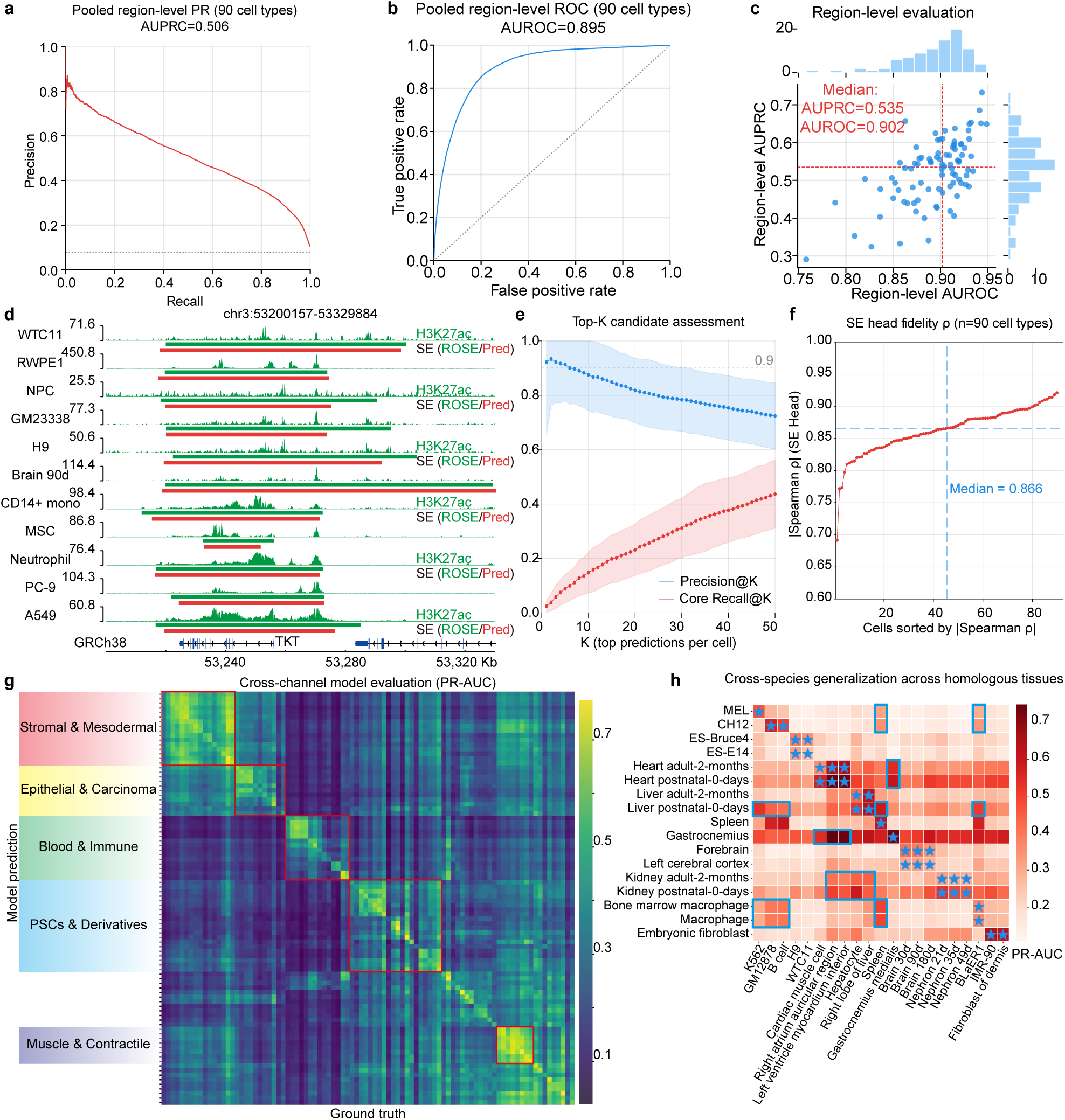
Genome-wide performance evaluation, in silico prioritization, and lineage-specific fidelity of Borzoi-SE. **a**, Overall region-level precision–recall (PR) curve on held-out test chromosomes (Split 0: chr3 and chr11). **b**, Overall region-level receiver operating characteristic (ROC) curve on held-out test chromosomes (Split 0: chr3 and chr11). **c**, Distribution of region-level AUROC (*x*-axis) and AUPRC (*y*-axis) across all *n* = 90 modeled human cell types and tissues, with marginal density histograms. **d**, Representative genomic snapshot at the *TKT* locus (chr3:53,200,157–53,329,884; ∼130 kb) across 11 representative lineages, displaying normalized experimental H3K27ac ChIP-seq coverage (top track), ROSE ground-truth SEs (green), and Borzoi-SE-predicted SE probabilities (red). **e**, *In silico* candidate prioritization curves across top *K* predictions (*K* = 1 to 50): Precision@K (blue) and Core_Recall@K (red; recovery rate of the top 50% highest H3K27ac-ranked SEs). Solid lines and shaded bands denote mean ± s.d. across *n* = 90 cell types. **f**, Absolute Spearman correlations between summed predicted SE probabilities and empirical ROSE ranks across 90 cell types, ordered by correlation magnitude (median |ρ| = 0.866).. **g**, Pairwise cross-channel evaluation matrix (AUPRC) comparing Borzoi-SE predictions (rows, grouped by major physiological clades) against experimental ground truth (columns) across all 90 cell types. **h**, Cross-species zero-shot generalization heatmap (AUPRC) evaluating human prediction channels (columns) directly on mouse genomic sequences against empirical mouse SE annotations (rows). Blue stars indicate matched human–mouse cell or tissue pairs, and blue boxes highlight off-diagonal cross-predictions consistent with developmental relationships between tissues.

To evaluate Borzoi-SE’s capacity for *in silico* prioritization, we applied the validation-selected decision threshold to the held-out test chromosomes, which yielded balanced global performance (precision ≈ 0.55, recall ≈ 0.40) and maintained consistent predictive accuracy regardless of the variable super-enhancer abundance across distinct cell types (**Supplementary Fig. 2a**). At the locus level, Borzoi-SE faithfully recapitulated lineage-restricted chromatin activation patterns. For example, at the ∼130-kb *TKT* super-enhancer locus on chromosome 3, the model correctly predicted epigenomic quiescence in non-expressing lineages (such as WTC11 and RWPE1) while robustly identifying active super-enhancers across responsive cellular contexts, precisely mirroring experimental H3K27ac occupancy (**Fig. 2d**).

To simulate prospective experimental prioritization workflows, we performed a threshold-free top-*K* candidate ranking based on prediction confidence. Borzoi-SE maintained high validation precision across escalating rank depths (precision of approximately 0.90 at K = 1 and above 0.70 at K = 50), demonstrating stringent false-discovery control (**Fig. 2e**). Furthermore, when evaluating the capacity of Borzoi-SE to identify core regulatory hubs, defined as the top 50% most active super-enhancers by empirical H3K27ac intensity, these dominant elements were preferentially enriched among the highest-confidence predictions, achieving a core recall exceeding 0.45 at *K* = 50 (**Fig. 2e**). These results demonstrate that Borzoi-SE inherently internalizes the quantitative hierarchy of cis-regulatory elements without requiring explicit signal strength supervision. Although Borzoi-SE demonstrated robust precision across both global evaluations and top-*K* prioritization, its overall recall remained more moderate (global recall ≈ 0.40; **Supplementary Fig. 2a**). To investigate whether this asymmetry reflects stochastic omission or is governed by intrinsic biological properties, we stratified detection recall across deciles of experimental super-enhancer strength (**Supplementary Fig. 2b**). Detection recall exhibited an overall monotonic decline from the strongest to the weakest elements (from 0.61 ± 0.20 in Decile 1 to 0.23 ± 0.14 in Decile 10; **Supplementary Fig. 2b**). This indicates that false negatives are predominantly confined to borderline elements near the empirical ROSE threshold, whereas dominant, high-potency regulatory hubs are captured with high sensitivity.

Because detection sensitivity was strictly coupled to biological strength, we next asked whether Borzoi-SE quantitatively decodes the continuous potency of cis-regulatory elements. We defined an *in silico* proxy for enhancer activity by summing predicted SE probabilities across stitched enhancers and aligned these scores against experimental ROSE rankings (**Fig. 2f**). Across all 90 cell types, predicted probability sums correlated strongly with biological SE ranks (absolute Spearman |*ρ*| = 0.69–0.92, median 0.866 ; **Fig. 2f**), with representative lineages demonstrating that super-enhancers receive disproportionately higher predicted scores than typical enhancers (**Supplementary Fig. 2c**). Together, these findings establish that Borzoi-SE internalizes the continuous regulatory hierarchy that distinguishes true super-enhancers from typical enhancers, prioritizing functionally dominant elements from sequence alone without requiring explicit signal strength supervision.

To determine whether Borzoi-SE relies on authentic lineage-specific syntax rather than ubiquitous chromatin features, we cross-evaluated prediction channels against all pairwise cell-type ground truths (**Fig. 2g; Supplementary Fig. 2d**). This pairwise evaluation revealed marked lineage fidelity: predictive accuracy peaked sharply along the matched diagonal (concordant PR-AUC > 0.5–0.7; ROC-AUC > 0.90–0.95) and collapsed between unrelated cell types (off-diagonal PR-AUC < 0.1–0.2). Moderate off-diagonal concordance was mainly observed among related cell types, forming distinct lineage groups such as hematopoietic/immune, pluripotent, muscle, epithelial, and stromal cells (red boxes; **Fig. 2g**). These related cell types likely share similar super-enhancer landscapes because they are controlled by overlapping sets of master transcription factors. This modular concordance demonstrates that Borzoi-SE decodes genuine cell-type-specific regulatory grammars rather than non-specific open-chromatin motifs.

We next evaluated whether this internalized syntax generalizes across mammalian evolution by applying human prediction channels directly to mouse genomic sequences from matched tissues in a zero-shot setting (**Fig. 2h**). Functionally homologous tissue pairs (e.g., embryonic stem cells, liver, heart, and blood/spleen; blue stars) exhibited high predictive concordance with mouse SEs derived from empirical H3K27ac profiles (PR-AUC > 0.5–0.7; **Supplementary Table 3**). Importantly, prominent off-diagonal cross-predictions (blue boxes; **Fig. 2h**) reflected developmental transitions rather than technical noise. For instance, mouse postnatal day 0 liver strongly cross-predicted human hematopoietic and lymphoid channels (K562, GM12878, B cells, spleen, and BLaER1), recapitulating the physiological role of the perinatal mammalian liver as a primary site of extramedullary hematopoiesis prior to marrow colonization^18^.

Together, these findings establish that Borzoi-SE internalizes conserved sequence grammar governing cell identity across mammalian evolution, setting the stage for base-resolution mechanistic dissection of its driving regulatory features.

### Borzoi-SE resolves base-resolution cis-regulatory grammar and lineage-specific master TF syntax

To determine whether Borzoi-SE relies on authentic biological sequence syntax, we applied a gradient-weighted input (Gradient × Input) attribution framework to quantify base-resolution nucleotide contribution scores across input sequences. We first evaluated this interpretability framework at the classical human *β*-globin locus control region (LCR) on chromosome 11 (chr11:5,223,000–5,304,000; ∼81 kb) in K562 erythroleukemia cells (**Fig. 3a**). Borzoi-SE accurately predicted the super-enhancer landscape, directly co-localizing with ground-truth ROSE-defined SE boundaries and marked H3K27ac accumulation. This predicted domain also aligned with local chromatin accessibility (ATAC-seq) and established binding profiles of master erythroid TFs (GATA1 and NFE2) and the structural organizer CTCF (**Fig. 3a**). Base-resolution attribution across the five constituent hypersensitive sites (HS1–HS5) resolved the underlying regulatory syntax: HS1–HS4 were enriched for motifs of erythroid TFs, including GATA1::TAL1, KLF1 and MAF::NFE2 motifs, while HS5 was dominated by the canonical CTCF insulator motif (**Fig. 3b**). Thus, without explicit ChIP-seq supervision, Borzoi-SE resolves both the core master-factor binding sites and chromatin boundary insulation governing in vivo super-enhancer organization *de novo*.

**Figure 3:**
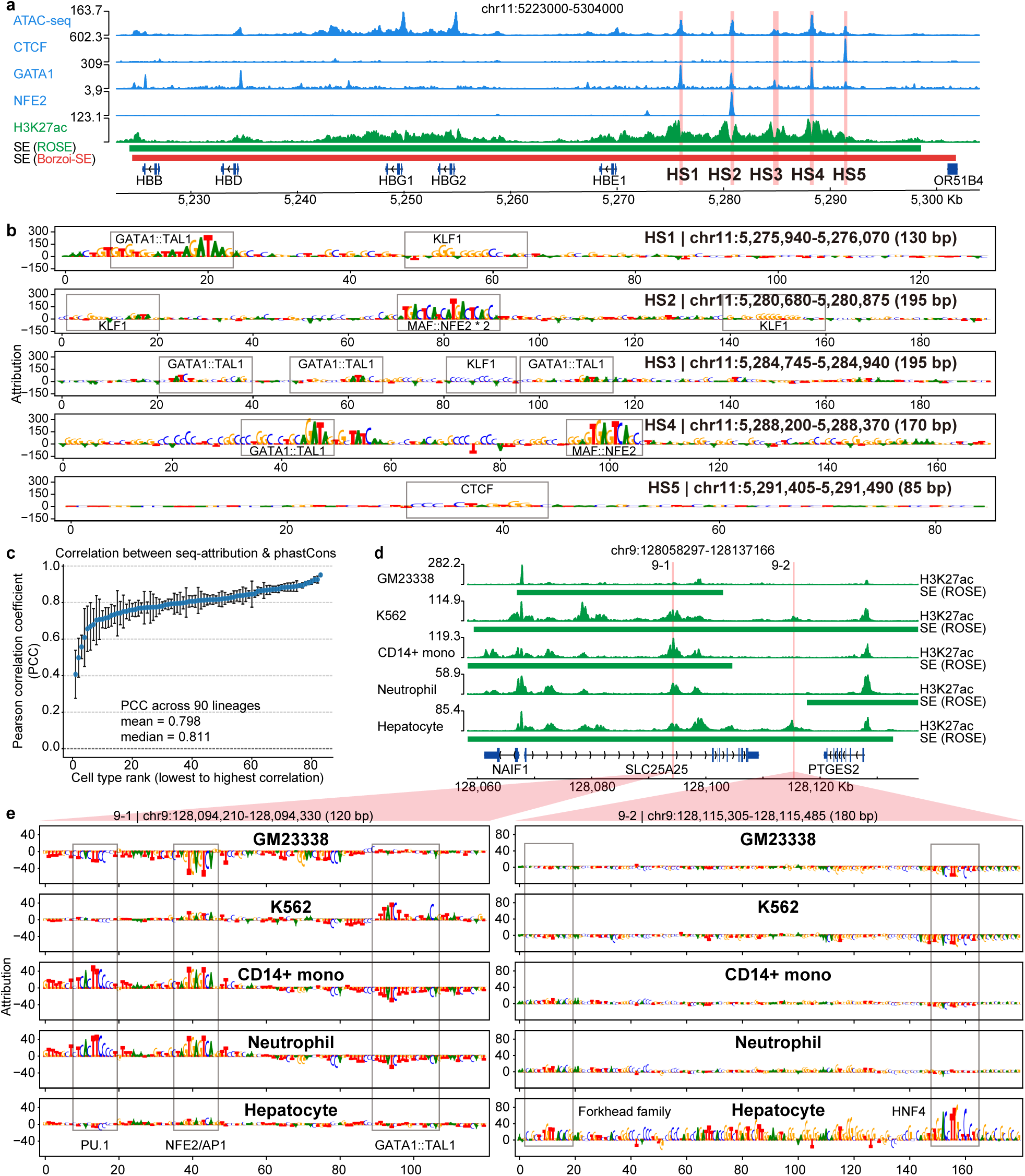
Borzoi-SE decodes base-resolution cis-regulatory grammar, evolutionary conservation, and lineage-specific master transcription factor switching. **a**, Genomic snapshot of the human *β*-globin locus control region (LCR; chr11:5,223,000–5,304,000; ∼81 kb) in K562 cells, displaying normalized ATAC-seq, CTCF, GATA1, NFE2, and H3K27ac tracks alongside ground-truth ROSE SEs (green) and Borzoi-SE-predicted SE probabilities (red). Classical constituents HS1–HS5 are highlighted by red vertical bars. **b**, Base-resolution Gradient × Input sequence attribution tracks across LCR constituents HS1–HS5 in K562 cells, annotating key TF motifs (GATA1::TAL1, MAF::NFE2, KLF1, and the structural insulator CTCF). **c**, Correlation between Borzoi-SE sequence attribution scores and vertebrate evolutionary conservation (phastCons 100-way) across *n* = 90 modeled human lineages. Points represent the mean Pearson correlation coefficient (*r*) for each cell type; error bars denote ± s.e.m. **d**, Multi-lineage genomic snapshot across the *SLC25A25* super-enhancer locus (chr9:128,058,297– 128,137,166; ∼78.9 kb) in five representative lineages, showing normalized H3K27ac ChIP-seq signal and ROSE SEs. Shaded vertical stripes mark constituent enhancers 9-1 and 9-2. **e**, Base-resolution DeepLIFT sequence attribution profiles across the five cell types at constituent enhancers 9-1 and 9-2.

We next asked whether this sequence prioritization is globally coupled to evolutionary constraint across the human genome. To reduce single-nucleotide attribution noise, we implemented a decile-stratified correlation analysis between binned attribution magnitude (10-bp sliding windows, 5-bp step) and vertebrate evolutionary conservation scores (PhastCons 100-way; **Methods**). Across all 90 modeled cell types, binned model attribution strongly correlated with evolutionary conservation (**Fig. 3c**). This genome-wide concordance confirms that the sequence features prioritized by Borzoi-SE are constrained by purifying selection across mammalian evolution.

Because super-enhancers are primarily orchestrated by lineage-defining master regulators, we tested whether coupling Gradient × Input profiles with the TF-MoDISco algorithm could recover these trans-acting factors *de novo*^19^. Unbiased motif extraction across cellular lineages successfully identified canonical lineage-specifying transcriptional circuits (**Supplementary Fig. 3a**): the OCT4/SOX2/NANOG pluripotency network in WTC11 and H9 stem cells, erythroid drivers (GATA1/NFE2/KLF) in K562, myeloid regulators (PU.1/CEBP/BATF) in CD14⁺ monocytes, and hepatic factors (HNF4/FOXA) in primary hepatocytes. These extractions confirm that the model identifies the specific TF repertoires operating within each lineage directly from DNA sequences.

Finally, we investigated how Borzoi-SE resolves super-enhancer domains that appear broadly active across multiple tissues at the epigenomic level. At the *SLC25A25* locus (∼78.9 kb), the model recapitulated H3K27ac-defined domain activity across diverse cell lineages (**Fig. 3d**). However, base-resolution attribution across two constituent enhancer hubs (sites 9-1 and 9-2) revealed distinct regulatory strategies (**Fig. 3e**). Whereas site 9-2 displayed a lineage-specific contribution, site 9-1 was active across both erythroid and myeloid contexts. Notably, at site 9-1, Borzoi-SE dynamically shifted its sequence attention to distinct motifs across cell types: in erythroid K562 cells, attribution concentrated on GATA1::TAL1, whereas in myeloid CD14⁺ monocytes and neutrophils, attribution relocated to PU.1 and NFE2/AP-1 motifs (**Fig. 3e**). A similar modular constituent organization was independently observed at the *ZMYND8* super-enhancer domain on chromosome 20 (**Supplementary Fig. 3b, c**), suggesting that a single CRE could harbor distinct motif sets that are used in different lineages.

This lineage-dependent reallocation of sequence attribution demonstrates that Borzoi-SE focuses on the functional drivers of each cellular context, allocating sequence attention specifically to the lineage-specific TFs operating in that cell state. This capability explains how broadly shared regulatory domains are assembled from modular, cell-type-specific cis-regulatory syntax.

### In silico constituent ablation and CRISPRi validation establish HS2 as the dominant hub of the β-globin LCR

Having decoded the localized sequence syntax of individual constituent elements, we next asked whether Borzoi-SE can resolve the higher-order constituent cooperativity and functional hierarchies that govern the assembly of intact, multi-constituent super-enhancer domains. Constituents of SEs often display non-additive, synergistic regulatory behavior, wherein specific “hub” or “facilitator” elements orchestrate the transcriptional activity of the entire domain^20–23^. To evaluate whether Borzoi-SE captures this higher-order hierarchy, we deployed the model to perform systematic *in silico* mutagenesis across two classical erythroid paradigms: the human *α*-globin and *β*-globin locus control regions (LCRs, here analyzed as SE domains). By computationally ablating individual constituent enhancers, promoters, and boundary elements, we quantified the resulting change in predicted super-enhancer probability (Δ = SE_prob_MUT_ − SE_prob_WT_) and calculated the relative locus activity integral ratio (Integral_MUT_/ Integral_WT_) in erythroleukemic K562 cells (**Fig. 4a-d**).

**Figure 4:**
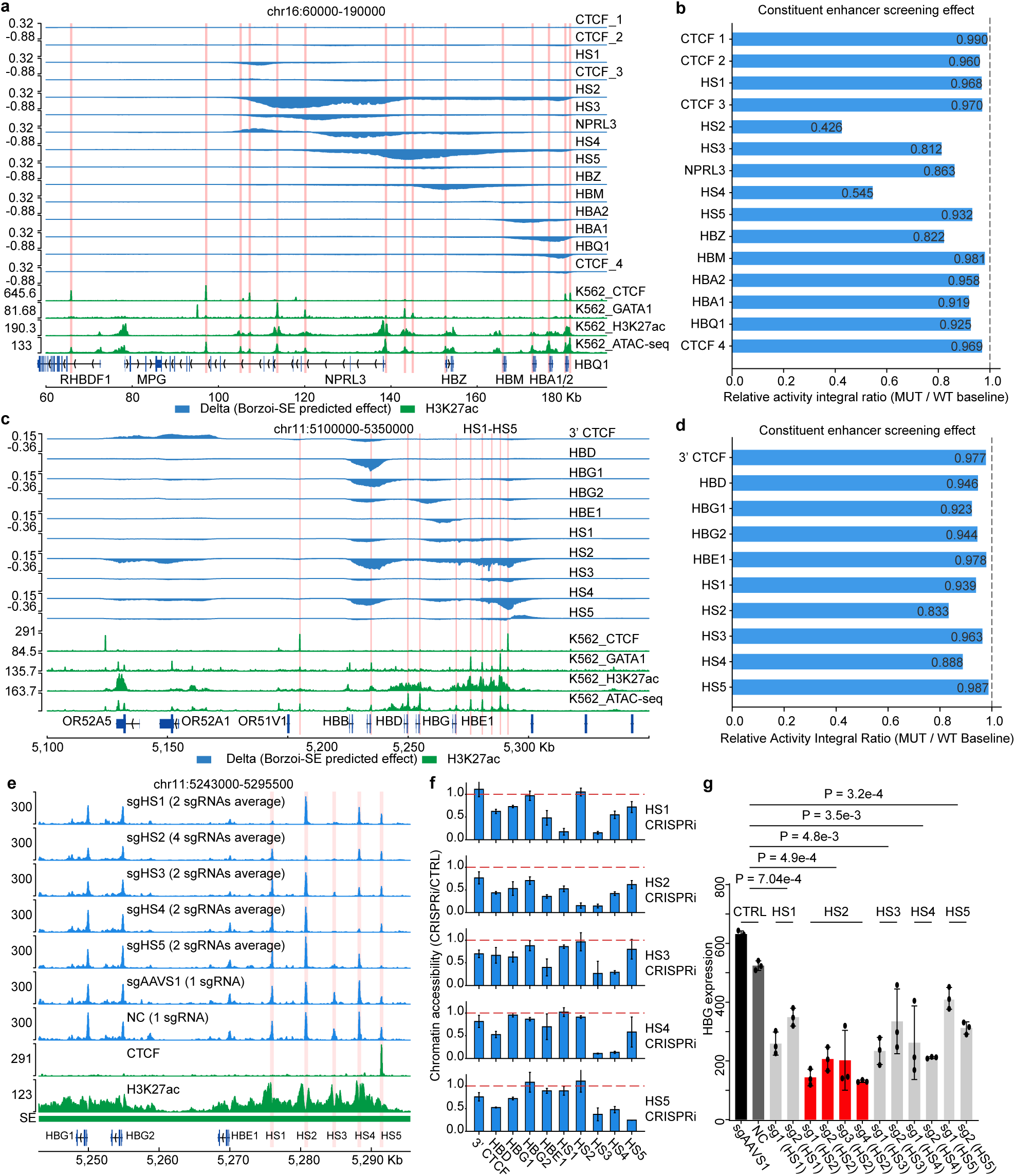
Borzoi-SE decodes constituent enhancer hierarchy and synergistic epistasis, validated by CRISPRi. **a**, *In silico* perturbation snapshot across the human *α*-globin LCR (chr16:60,000–190,000) in K562 cells. Top, predicted change in super-enhancer probability (Δ = SE_prob_MUT_ − SE_prob_WT_, dark blue) following *in silico* ablation of individual constituents. Bottom, normalized baseline experimental coverage for CTCF, GATA1, H3K27ac, and ATAC-seq. Red shaded bars indicate perturbation sites. **b**, Relative super-enhancer activity integral ratios across the constituent elements at the *α*-globin locus relative to the wild-type baseline (dashed line). **c**, *In silico* perturbation snapshot across the human *β*-globin LCR (chr11:5,100,000–5,350,000) in K562 cells. Tracks are arranged as in **a**, showing 10 *in silico* ablations of LCR constituents alongside baseline epigenomic profiles. **d**, Relative super-enhancer activity integral ratios across the 10 screened regulatory elements at the *β*-globin locus. **e**, Chromatin accessibility profiling across the *β*-globin LCR in K562 cells, showing normalized ATAC-seq tracks following CRISPRi targeting each LCR constituent alongside non-targeting control (NC) and negative genomic control (sgAAVS1). The signals of each track represent the average ATAC-seq signals for 1, 2 or 4 sgRNAs as indicated. **f**, Relative chromatin accessibility in **e** across 10 regulatory sites, normalized to negative controls (mean of NC and sgAAVS1; red dashed line). Error bars denote 95% CI (Student’s *t*-distribution across biological replicates). **g**, RT-qPCR quantification of *HBG* (*HBG1* and *HBG2*) expression following CRISPRi of constituent enhancers. 2-4 sgRNAs targeting each enhancer were used in parallel. Data are mean ± s.d. from 3 technical replicates; statistical significance was assessed using a two-tailed Student’s *t*-test.

Across the 15 regulatory elements screened within the ∼ 130-kb *HBA* LCR (chr16:60,000– 190,000), computational ablation of HS2 and HS4 precipitated the most severe collapse of cumulative super-enhancer activity, yielding activity integral ratios of 0.426 and 0.545, respectively (**Fig. 4a, b**). In contrast, ablations of intervening globin gene promoters and secondary hypersensitive sites produced only modest reductions (ratios: 0.812–0.981), while flanking CTCF boundaries showed negligible impact on core domain potency (ratios: 0.960–0.990; **Fig. 4a, b**). Notably, this computationally derived prioritization independently aligns with recent functional dissections of the orthologous mouse *α*-globin super-enhancer, where synthetic reconstitution demonstrated that the multispecies conserved sequence (MCS) element MCS-R2 (homologous to HS2 in K562) serves as the primary classical enhancer whose loss most severely impairs locus activation, operating alongside secondary facilitator elements including MCS-R4 (homologous to HS4 in K562)^20,21,24^. This cross-species concordance confirms that Borzoi-SE identifies authentic, evolutionarily conserved functional hubs within compound regulatory domains from primary sequence alone.

An unbiased *in silico* screen across 10 elements within the 250-kb *β* -globin locus (chr11:5,100,000–5,350,000) similarly identified HS2 as the preeminent functional hub of the *β*-globin LCR (**Fig. 4c, d**). Ablation of HS2 produced the strongest locus-wide suppression, reducing the total activity integral ratio to 0.833 and triggering broad probability depletion across the entire LCR domain and downstream promoters (**Fig. 4c, d**). In contrast, ablating individual promoters (such as *HBE1*, ratio = 0.978) or secondary hypersensitive sites (HS1, HS3, HS4, HS5) caused strictly focal perturbations (ratios: 0.888–0.987; **Fig. 4c, d**). Furthermore, computational ablation of the structural boundary elements, specifically the 3’ flanking CTCF (3’ CTCF) and the insulator HS5, triggered localized positive probability deltas in flanking regions, precisely modeling the boundary leakage and ectopic super-enhancer expansion that occurs upon the loss of CTCF insulation^25,26^ (**Fig. 4c**). Base-resolution attribution linked this hierarchical dominance directly to the sequence syntax of HS2, which harbors a high-affinity GATA1::TAL1 composite motif flanked by tandem MAF::NFE2 elements that supply the predominant positive weights across the locus^27–29^ (**Fig. 3b**).

To experimentally validate these *in silico* hierarchies, we carried out CRISPR interference (CRISPRi) against each constituent of the *β* -globin LCR (HS1 through HS5) in K562 cells, and measured chromatin accessibility by ATAC-seq (**Fig. 4e, f**) and target gene expression by RT-qPCR (**Fig. 4g**). Consistent with model predictions, targeted silencing of HS1, HS3, HS4, or HS5 induced strictly localized chromatin closure at their respective target sites (relative accessibility < 0.10–0.20), leaving neighboring hypersensitive sites and globin promoters largely intact (accessibility: 0.60–1.00; **Fig. 4e, f**). In contrast, transcriptional silencing of HS2 (sgHS2) triggered a collapse of chromatin accessibility across the entire LCR and its downstream target promoters, reducing relative accessibility at *HBD*, *HBG1*, *HBG2*, *HBE1*, HS1, HS3, and HS4 by 35% to 85% (**Fig. 4e, f**). At the transcriptional level, this structural collapse manifested as a severe, coordinated downregulation of fetal *β*-like globin genes (*HBG1/2*) specifically in sgHS2-treated cells, whereas targeting other individual hypersensitive sites led to only modest downregulation (**Fig. 4g**).

These findings establish that Borzoi-SE reliably captures constituent hierarchy within SEs.

### Borzoi-SE accurately predicts cell-type-specific pathogenic gain-of-SE mutations in T-ALL

Pathogenic non-coding mutations that generate *de novo* super-enhancers (Gain-of-SE) represent a critical class of oncogenic drivers that are exceptionally challenging to predict prospectively, as traditional epigenomic profiling cannot anticipate *de novo* chromatin emergence prior to empirical observation. To determine whether Borzoi-SE can predict such gain-of-function events ab initio without relying on pre-existing chromatin marks, we modeled a classic non-coding mutation upstream of the *TAL1* oncogene. This recurrent insertion creates a *de novo* binding motif for the MYB transcription factor, nucleating a downstream super-enhancer that drives aberrant *TAL1* overexpression in T-cell acute lymphoblastic leukemia (T-ALL)^16^.

We performed in silico mutagenesis by introducing a 12-bp insertion (CCGTTTCCTAAC at chr1:47,239,295) that creates *de novo* MYB binding motifs into the reference sequence, calculating the per-bin difference in predicted super-enhancer probability (Δ = SE_prob_MUT_ − SE_prob_WT_) across all 90 cell types (**Fig. 5a, c**)^16^. Borzoi-SE predicted a sharp, localized gain in super-enhancer probability directly upstream of the *TAL1* promoter that was strictly confined to hematopoietic lineages (**Fig. 5a**). Across all 90 evaluated channels, the top-ranked responses occupied the extreme right tail of the effect-size distribution, led by T-ALL models (KOPT-K1, Δ = 0.232; DND-41, Δ = 0.102), erythroleukemia (K562, Δ = 0.201), spleen (Δ = 0.121), HL-60 (Δ = 0.114), and BLaER1 (Δ = 0.092) (**Fig. 5b**). In contrast, non-hematopoietic control tissues exhibited completely flat response profiles (**Fig. 5a, b**).

**Figure 5:**
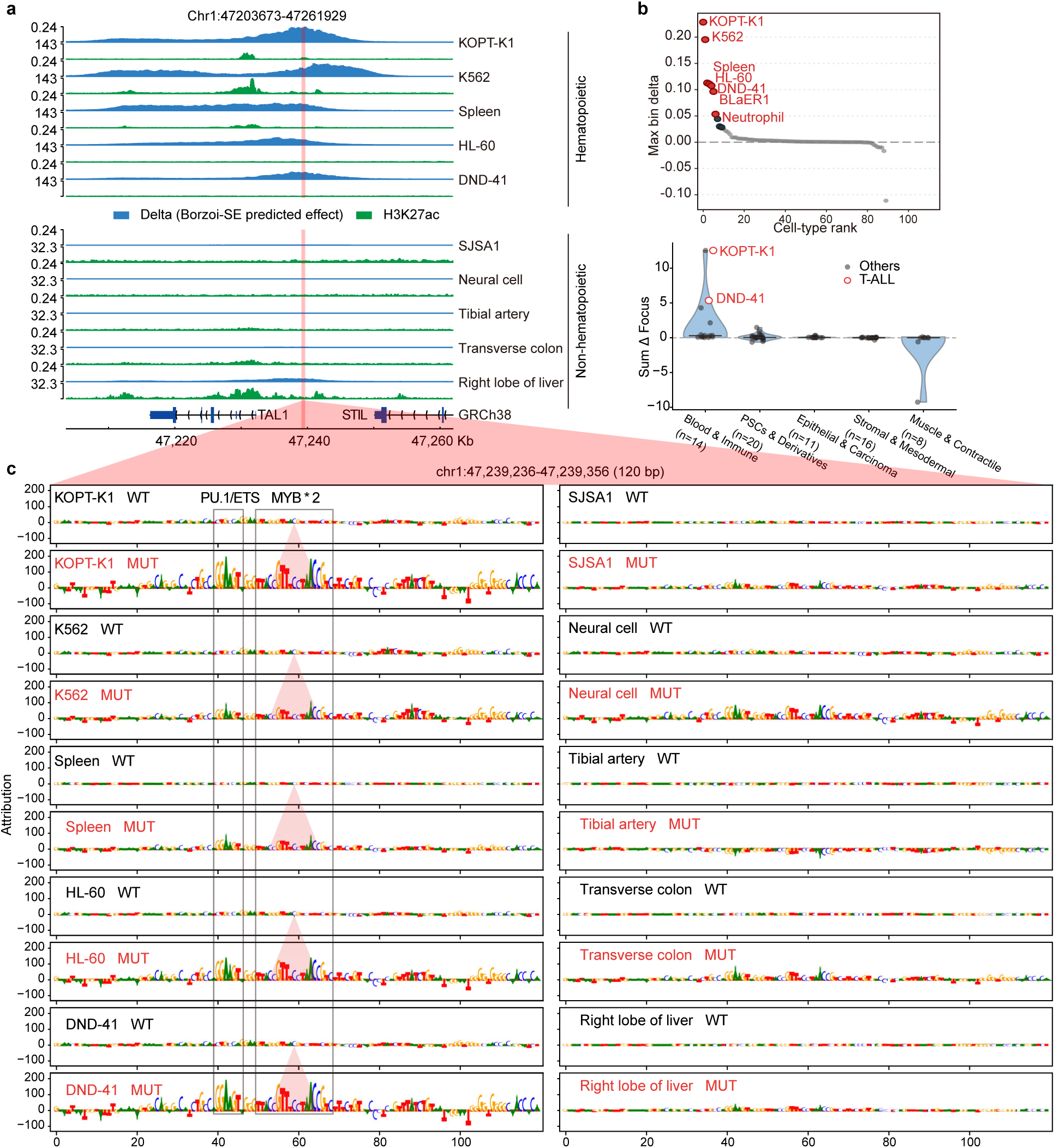
Borzoi-SE accurately predicts cell-type-specific pathogenic gain-of-super-enhancer variants in T-cell acute lymphoblastic leukemia (T-ALL). **a**, Multi-lineage genomic snapshot across the human *TAL1* locus (chr1:47,203,673–47,261,929). Top tracks depict predicted super-enhancer probability changes (Δ = SE_prob_MUT_ − SE_prob_WT_); bottom tracks display baseline experimental H3K27ac ChIP-seq coverage in wild-type cells. Lineages are stratified into responsive hematopoietic/leukemic lineages (top five) and non-responsive control lineages (bottom five). **b**, Top, rank-ordered distribution of maximum per-bin super-enhancer probability changes across *n* = 90 human cell types, highlighting responsive hematopoietic lineages in red. Bottom, violin plots of cumulative focal probability changes (Sum Δ Focus) stratified across five major physiological clades (defined in Fig. 2g). **c**, Base-resolution DeepLIFT sequence attribution profiles across a 120-bp window spanning wild-type and mutant alleles, comparing five responsive hematopoietic/leukemic lineages (left) and five non-responsive control lineages (right).

Because K562 erythroleukemia cells natively harbor strong H3K27ac accumulation at the wild-type locus, we investigated whether these predicted gains merely reflected the memory of baseline epigenetic status. Notably, in T-ALL lines (KOPT-K1 and DND-41) and other responsive lineages, the wild-type locus completely lacked baseline H3K27ac enrichment, yet these models exhibited the highest *de novo* induction scores (**Fig. 5a**). Resolving sequence importance via DeepLIFT attribution across a 120-bp centered window confirmed the underlying mechanistic syntax: the mutant allele selectively acquired high positive attribution scores at the newly generated MYB motif strictly within responsive leukemic channels (attribution peaks > 150 in KOPT-K1 and DND-41) (**Fig. 5c**)^30^. Crucially, beyond activating the insertion site itself, this *de novo* motif cooperatively awakened surrounding cis-regulatory grammar, substantially elevating the sequence attribution of flanking endogenous motifs, such as an adjacent upstream PU.1/ETS site (∼40–45 bp), that remained dormant in the wild-type sequence; in contrast, non-responsive control tissues maintained flat attribution across both alleles **(Fig. 5c)**.

Having confirmed that Borzoi-SE resolves de novo domain nucleation driven by a focal 12-bp insertion at a paradigmatic oncogenic locus, we next evaluated whether single-nucleotide polymorphisms (SNPs) can quantitatively alter super-enhancer homeostasis across the human genome. By scaling in silico mutagenesis genome-wide, we aimed to systematically prioritize both activating and repressive variants independent of linkage disequilibrium.

### Genome-wide in silico screening prioritizes functional non-coding variants calibrated by population haQTLs

Dissecting the functional mechanisms of non-coding genetic variants identified by GWAS and QTL mapping remains a formidable challenge. Many post-GWAS workflows prioritize variants by their overlap with annotated cis-regulatory elements (CREs). Although useful for identifying variants that alter existing CREs, this approach can miss gain-of-function variants that establish de novo enhancer activity in regions lacking regulatory annotation^31–34^. Because Borzoi-SE learns intrinsic sequence-to-function grammar independent of prior functional annotations, it provides a powerful computational framework to prioritize both activating and repressive candidate regulatory variants directly from sequence^15^.

To evaluate Borzoi-SE in functional variant screening, we modeled genetic variants across diverse output channels capturing multiple sub-lineages and developmental stages within the brain, heart, and muscle systems. For each variant, we computed the change in predicted super-enhancer probability (Δ = SE_prob_MUT_ − SE_prob_WT_) across 256-bp bins and categorized variants into neutral, candidate GoF (predicted activation from a low baseline probability), and candidate LoF (predicted attenuation of regulatory potential) classes. Across all systems, variants with substantial predicted regulatory impact (∣ Δ ∣≥ 0.025) were exceptionally sparse, confirming stringent specificity against neutral background variation (**Fig. 6a**). When surveyed across the full compendium of developmental and lineage-specific channels, prioritized candidates exhibited prominent cell-state selectivity across physiological subtypes (**Fig. 6b**). Next, we asked whether this in silico scoring can resolve a central dilemma in GWAS fine-mapping: distinguishing true causal drivers from non-functional linkage disequilibrium (LD) passenger variants^35^.

**Figure 6:**
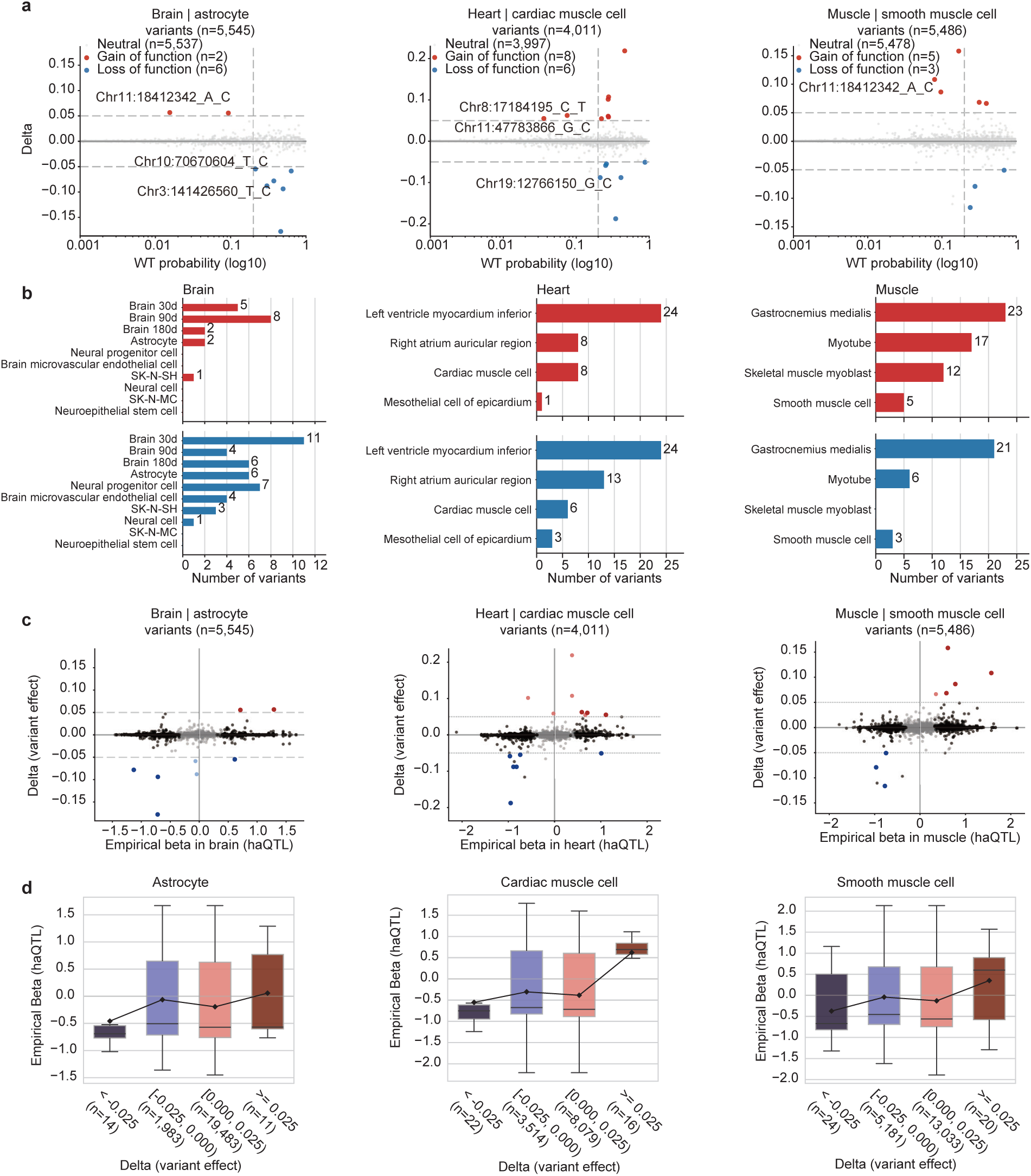
Borzoi-SE conducts *ab initio* fine-mapping to identify functional variants and *de novo* gain-of-function super-enhancers validated by population haQTLs. **a**, Genome-wide *in silico* variant prioritization and functional classification across brain, heart, and muscle systems. Scatter plots show predicted super-enhancer probability change (Δ = SE_prob_MUT_ − SE_prob_WT_) versus wild-type baseline probability, partitioning variants into neutral (grey/black), gain-of-function (GoF; red), and loss-of-function (LoF; blue). **b**, Number of prioritized functional GoF (red) and LoF (blue) variants stratified across specific cell types and developmental stages within brain, heart, and muscle lineages. **c**, Concordance between *in silico* variant effect predictions (Δ) and empirical histone acetylation QTL (haQTL) effect sizes (*β*) across astrocytes (brain), cardiac muscle cells (heart), and smooth muscle cells (muscle). **d**, Distribution of empirical haQTL effect sizes (*β*) across four stratified tiers of predicted impact (Δ < −0.025, [−0.025,0.000), [0.000,0.025), and Δ ≥ 0.025) in astrocytes, cardiac muscle cells, and smooth muscle cells. Center lines denote medians; box limits indicate interquartile range (IQR); whiskers show non-outlier ranges; black diamonds and connecting lines indicate bin means.

We next benchmarked model-predicted Δ scores against empirical histone acetylation quantitative trait locus (haQTL) effect sizes derived from matched GTEx tissues^36^. In population genetics, directly correlating computational sequence scores with raw haQTL effect sizes frequently yields weak associations. This attenuation arises because statistical QTL mapping is heavily confounded by linkage disequilibrium (LD), which assigns significant effect sizes to non-causal “passenger” variants co-inherited within the same haplotype block^12^. By evaluating the intrinsic sequence consequence of individual nucleotide substitutions, Borzoi-SE helps disentangle likely functional drivers from LD hitchhikers. In the unfiltered pool, the vast majority of variants yielded negligible predicted sequence impact (Δ ≈ 0) despite displaying a wide spectrum of non-zero empirical effect sizes (haQTL Beta), a classic hallmark of non-functional LD passenger variants. In contrast, variants prioritized by Borzoi-SE with substantial predicted impacts displayed a robust positive concordance with empirical haQTL effect sizes across all three physiological systems (**Fig. 6c**). To quantify this concordance, we categorized variants into escalating effect-size groups. Across all three tissue systems, empirical haQTL effect sizes exhibited a monotonic upward progression matching the model-predicted directionality (**Fig. 6d**). Together, these results demonstrate that Borzoi-SE provides effective *in silico* prioritization of functional regulatory variants, with predicted effect magnitudes quantitatively calibrating biological chromatin shifts. While this genome-wide screen effectively prioritizes variants with bidirectional regulatory potential, computational Δ scores alone do not resolve the underlying physical mechanisms. A key question is whether these prioritized mutations truly alter chromatin regulation, either by disrupting important TF motifs in existing regulatory elements or by creating new motifs that can activate de novo super-enhancers..

### Base-resolution attribution decodes functional non-coding variants and validates *de novo* cardiac super-enhancer nucleation

To investigate the sequence mechanisms underlying candidate repressive variants, base-resolution DeepLIFT attribution was performed on prioritized loss-of-function (LoF) mutations alongside integration of population-scale epigenomic and transcriptomic cohorts (**Supplementary Fig. 4a-d**)^30^.

At the myogenic *LDHA* locus (∼137.2 kb), a candidate G>C transversion (chr11:18,369,690_G_C; rs10832923) selectively depleted predicted super-enhancer activity across myogenic and contractile lineages (Δ = −0.14 *to* − 0.11) while leaving non-myogenic tissues unaffected (**Supplementary Fig. 4a**). Nucleotide-level attribution across the region revealed the underlying sequence lesion: the G>C substitution directly disrupts the core STAT3 motif, precipitating a concomitant attenuation of sequence attribution at the adjacent upstream MEF2C master regulator motif (**Supplementary Fig. 4c**). This motif abrogation directly translated into concordant epigenetic, transcriptional, and organismal repression in human cohorts: in GTEx skeletal muscle, rs10832923 acts as a potent repressive haQTL (nominal *P* = 3.98 × 10^−7^, Beta = -0.815) and drives significant eQTL downregulation of *LDHA* (*P* = 7.1 × 10^−5^^7^, normalized effect size (NES) = -0.31), while fine-mapped GWAS credible sets link this variant to whole-body fat-free mass (*P* = 4.73 × 10^−7^) and basal metabolic rate (*P* = 9.97 × 10^−8^), concordant with the vital role of *LDHA*-mediated anaerobic glycolysis in muscle energy homeostasis (**Supplementary Fig. 4a**)^36–38^. Similarly, at the *MAP2K6* domain (∼409.6 kb), a prioritized G>A transition (chr17:69,480,329_G_A; rs9907575) induced marked, lineage-restricted loss of super-enhancer activity in gastrocnemius medialis (Δ = −0.25) and left ventricular myocardium (Δ = −0.18) (**Supplementary Fig. 4b**). Base-resolution attribution confirmed that this single-base alteration destroys a conserved composite element composed of contiguous androgen/glucocorticoid receptor (AR/NR3C1) and nuclear factor I (NFI) motifs (**Supplementary Fig. 4d**). Corroborating this computational disruption, rs9907575 functions as a significant repressive haQTL in skeletal muscle (nominal *P* = 1.24 × 10^−9^, Beta = -0.910), is coupled to transcriptional downregulation of the adjacent kinase *MAP2K6* (*P* = 4.7 × 10^−22^, NES = -0.49), and associates with metabolic traits including increased waist-to-hip ratio (*P* = 7.00 × 10^−7^) and elevated circulating triglycerides (*P* = 8.18 × 10^−10^)^36–41^. Together, these analyses confirm that Borzoi-SE-prioritized LoF variants operate by dismantling essential master and cooperative transcription factor syntax, precipitating localized chromatin closure and target gene repression.

Having established that Borzoi-SE accurately models how single-nucleotide disruptions dismantle established regulatory elements, we turned to the converse and more challenging paradigm: identifying non-coding variants that introduce *de novo* regulatory grammar to nucleate super-enhancers within previously quiescent chromatin. Within our prioritized genome-wide screen, Borzoi-SE identified a non-coding T>C transition (chr7:136,884,669_T_C; rs1424569) at the *CHRM2* locus (∼409.6 kb) predicted to induce a massive, lineage-specific SE activation that was strictly confined to myocardium, whereas non-cardiac control tissues maintained flat baselines (**Fig. 7a**). Resolving nucleotide contribution via DeepLIFT attribution across a 100-bp centered window revealed the underlying *cis*-regulatory grammar: while the reference T allele exhibited quiescent baseline attribution, the mutant C allele creates a *de novo*, high-affinity androgen/glucocorticoid receptor (AR/NR3C1) consensus motif directly adjacent to an endogenous MEF2C binding site, which concomitantly boosted the sequence attribution of the neighboring MEF2C motif to establish a potent composite regulatory element (**Fig. 7b**)^30^.

**Figure 7:**
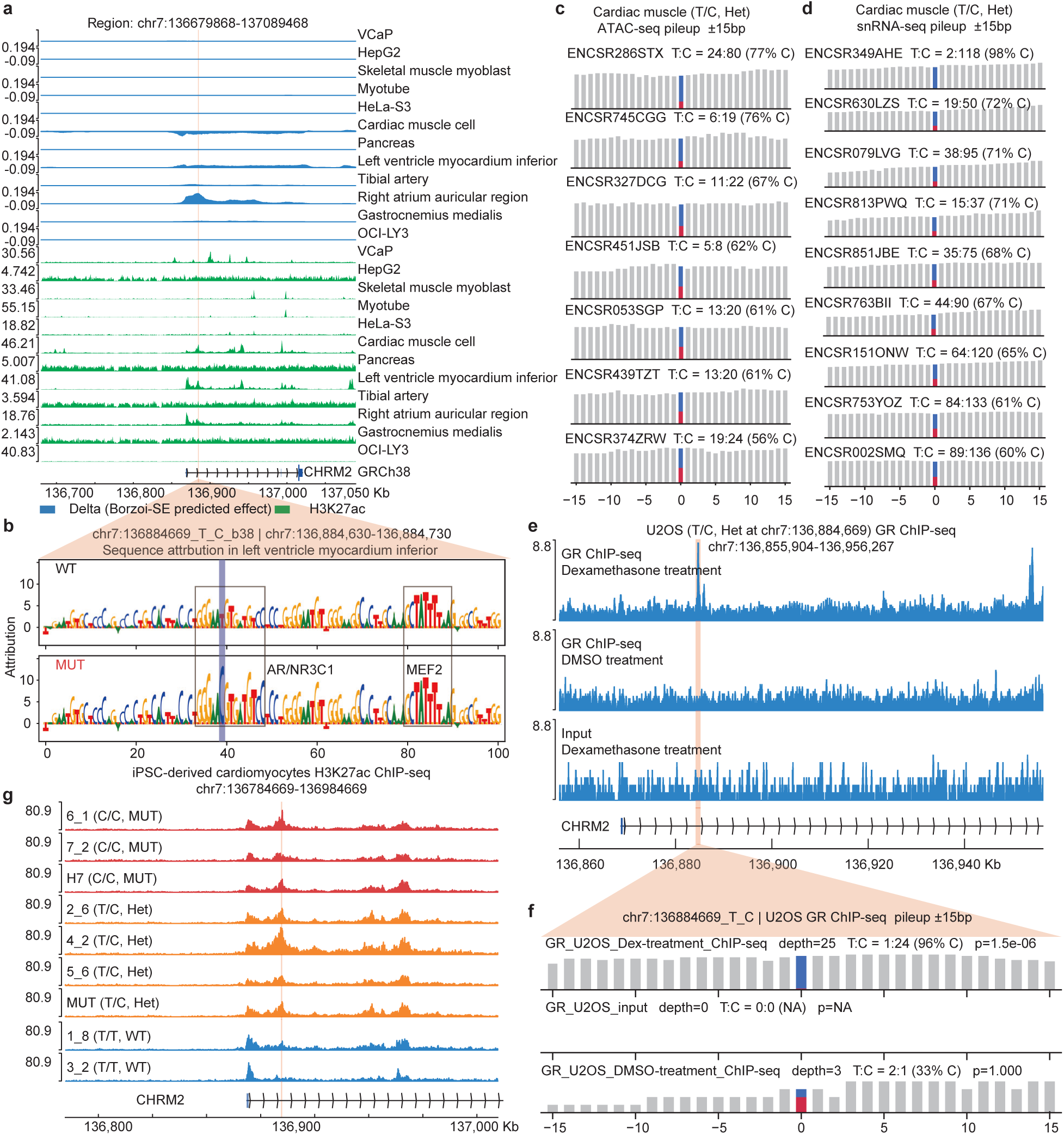
*De novo* gain-of-function super-enhancer discovery and multi-omics orthogonal validation across human cardiac traits. **a**, Multi-lineage genomic snapshot across the human *CHRM2* locus (chr7:136,679,868–137,089,468). Top tracks display predicted super-enhancer probability changes (Δ) following the T>C transition (chr7:136,884,669_T_C) across 12 human cell lines and primary tissues; bottom tracks display normalized baseline H3K27ac ChIP-seq coverage in wild-type cells. **b**, Base-resolution DeepLIFT sequence attribution profiles across chr7:136,884,630–136,884,730 comparing reference and alternative alleles in human left ventricular myocardium, highlighting the *de novo* creation of an AR/NR3C1 motif adjacent to an existing MEF2 motif. **c**, Allele-specific chromatin accessibility profiling via ATAC-seq read pileup (±15 bp) across *n* = 7 heterozygous human primary heart tissue samples from the ENCODE project. **d**, Allele-specific single-nucleus RNA-seq (snRNA-seq) read pileup (±15 bp) across *n* = 9 heterozygous human primary heart tissue samples from the ENCODE project. **e**, Genomic tracks across the *CHRM2* locus (chr7:136,855,904–136,956,267) depicting GR ChIP-seq coverage in naturally heterozygous human U2OS cells treated with dexamethasone (Dex), vehicle (DMSO), or input control. **f**, Allele-specific read pileup of GR ChIP-seq in heterozygous U2OS cells (± 15 bp around chr7:136,884,669_T_C). Under Dex treatment, 96% of captured reads originate from the mutant C allele (depth = 25; C = 24, T = 1). **g**, Normalized H3K27ac ChIP-seq coverage in iPSC-derived cardiomyocytes (iPSC-CMs; chr7:136,784,669–136,984,669) across homozygous mutant (red), heterozygous (orange), and wild-type (blue) lines. The red vertical line marks the mutation site (chr7:136,884,669_T_C).

This computationally predicted sequence activation matched empirical population-level genetics and organismal phenotypes. In the GWAS catalog, rs1424569 is significantly associated with prolongation of the electrocardiographic PR interval (GCST010321; *P* = 5.0 × 10^−8^, β = +0.36)^42^. Furthermore, fine-mapping of UK Biobank traits prioritized rs1424569 as the sole causal variant (posterior probability = 1.00; SuSiE-inf) for resting heart rate (*P* = 5.24 × 10^−17^, Beta = -0.0111)^38,43^. This finding is directly consistent with the established physiology of the encoded M2 muscarinic acetylcholine receptor (*CHRM2*), which slows atrioventricular nodal conduction to modulate cardiac rhythmicity^44^. In human cardiac tissues, rs1424569 acts as a significant histone acetylation quantitative trait locus (haQTL), driving local H3K27ac accumulation across the host *cis*-regulatory region (chr7:136,882,992–136,885,607; nominal *P* = 1.04 × 10^−4^, Beta = +0.567; and adjacent peak chr7:136,881,973–136,882,517; nominal *P* = 9.97 × 10^−8^, Beta = +0.898)^36^. Concordantly, GTEx analysis confirmed that this epigenetic activation drives robust transcriptional upregulation of *CHRM2* in human heart tissue (left ventricle: *P* = 8.6 × 10^−41^, NES = +0.28; atrial appendage: *P* = 7.2 × 10^−5^, NES = +0.072)^37^.

To track the physical molecular cascade initiated by this single-base mutation, we interrogated orthogonal epigenomic and transcriptional datasets across primary heart samples, genetically perturbed cellular models, and iPSC-derived cardiomyocytes (**Fig. 7c-g; Supplementary Table 5, Supplementary Fig. 5e**)^17,45,46^. After correction for reference-mapping bias using WASP (all WASP-corrected allele-specific analyses are detailed in **Supplementary Table 6**), allele-specific ATAC-seq in heterozygous primary human heart tissues revealed marked chromatin accessibility skewed toward the mutant C allele (62%–77% in left ventricle, 56%–67% in right ventricle; **Fig. 7c**)^47^. Single-nucleus RNA-seq (snRNA-seq) across nine heterozygous human heart samples confirmed that this chromatin opening directs allelic expression imbalance of *CHRM2*, with persistent skew toward the C allele (60%– 98% C; **Fig. 7d**). Direct transcription factor recruitment was verified in heterozygous U2OS cells treated with dexamethasone, a glucocorticoid agonist that triggers GR nuclear translocation^48^; GR ChIP-seq demonstrated that 96% of allele-informative reads localized to the mutant C allele (24 of 25 reads), whereas vehicle-treated controls yielded only background noise (**Fig. 7e, f)**^46^. In iPSC-derived cardiomyocytes, homozygous mutant and heterozygous lines exhibited elevated H3K27ac deposition across the mutation site compared to homozygous wild-type lines (**Fig. 7g**)^45^. Crucially, standard ROSE ranking demonstrated a status transition: while homozygous reference lines remained typical enhancers, several heterozygous lines crossed the geometric inflection point to form bona fide super-enhancers ranked 126–415 (**Supplementary Fig. 5e**).

A second GoF candidate, a C>A transversion (chr16:74,903,324_C_A; rs9937638) within the *WDR59* locus (∼409.6 kb; chr16:74,698,523–75,108,123), provided independent confirmation of this activation mode (**Supplementary Fig. 5a–e**). Borzoi-SE predicted a cardiac-restricted elevation in super-enhancer probability. DeepLIFT attribution confirmed that the mutant A allele creates a *de novo* GATA4 core recognition motif alongside an adjacent MEIS1/2 motif (**Supplementary Fig. 5b**)^30^. In matched cardiac tissue, rs9937638 mediates significant *cis*-regulatory H3K27ac enrichment within the surrounding regulatory domain (chr16:74,898,492–74,899,161; nominal *P* = 9.55 × 10^−6^, Beta = +0.840) and drives transcriptional upregulation of *WDR59* across cardiac compartments in GTEx (atrial appendage: *P* = 2.1 × 10^−50^, NES = +0.61; left ventricle: *P* = 1.8 × 10^−36^, NES = +0.47)^36,37^. In iPSC-derived cardiomyocytes, heterozygous and mutant lines exhibited marked accumulation of GATA4, H3K4me3, and H3K27ac (**Supplementary Table 5, Supplementary Fig. 5c**), accompanied by pronounced A-allele skews in primary heart ATAC-seq (56–67% A) and iPSC-derived cardiomyocyte GATA4 ChIP-seq (67%–94% A; **Supplementary Fig. 5d**), driving the progressive epigenomic activation of the locus (**Supplementary Fig. 5e**)^45^.

These findings demonstrate that Borzoi-SE identifies high-impact non-coding variants that introduce *de novo cis*-regulatory syntax capable of driving super-enhancer status transition. By capturing the physical chain of events from single-base motif creation and pioneer factor recruitment to domain-wide histone acetylation, target gene activation, and electrophysiological alterations, Borzoi-SE establishes an effective framework for decoding the non-coding genetic architecture of human disease.

## Discussion

The central conceptual advance of this study is that the defining properties of super-enhancers, including their domain-scale chromatin topology, continuous regulatory potency, and lineage-restricted activity, can be decoded directly from primary DNA sequence when the computational architecture matches the physical scale of genome organization. Existing sequence-based predictors frame super-enhancer discovery as binary classification of isolated, truncated snippets (≤ 3 kb), fundamentally decoupling enhancers from their spatial and cellular context^6–9^. In contrast, Borzoi-SE reads 524 kb of contiguous genomic sequence and outputs predictions across 1,600 spatial bins at 256-bp resolution across 90 human cell types, while decoupling constituent clustering (Cluster head) from super-enhancer discrimination (SE head, **Fig. 1a**). Each structural design directly corresponds to an underlying biological principle: the long-range input captures the broad ten-to-hundred-kilobase span of cooperative enhancer clusters; the spatial binning resolves precise boundary coordinates rather than assigning a single locus score; and the multi-lineage output channels preserve the dynamic, tissue-specific execution of master transcription factor (TF) programs. Consequently, Borzoi-SE captures higher-order regulatory syntax rather than localized motif density, as demonstrated by desert false-positive suppression and non-linear motif cooperativity.

Methodological rigor and experimental parity are paramount for contextualizing these advances. Rather than evaluating published models on historical, heterogeneously curated datasets, we established an unbiased benchmark wherein all four existing architectures (DeepSE, SENet, MuSE, and DcSE) were systematically retrained on our standardized H3K27ac-derived ROSE annotations under an identical, chromosome-isolated partitioning protocol^6–9^. Under this rigorous apples-to-apples evaluation, Borzoi-SE establishes decisive superiority across every evaluated dimension. In cluster-level binary classification, Borzoi-SE consistently outperforms baseline classifiers, confirming that foundation-scale representations and contextual priors enhance rather than dilute local motif discrimination. When deployed to continuous whole-chromosome screening, this architectural gap widens drastically: while fragment-based baselines suffer severe precision collapse due to dense false-positive clusters across non-coding gene deserts, Borzoi-SE suppresses off-target noise by approximately tenfold through its dilated Cluster head. Most notably, while baseline classifiers output single binary scalars lacking quantitative calibration, Borzoi-SE yields a cumulative spatial probability ( ∑ SE_prob ) that quantitatively scales with biological H3K27ac deposition and strongly correlates with empirical ROSE rankings (**Fig. 2f**). These convergent results confirm that long-range receptive fields and hierarchical object detection are essential for translating primary sequence into physiologically calibrated regulatory landscapes.

Beyond establishing benchmark-leading accuracy, nucleotide-resolution interpretability and functional perturbations provide direct mechanistic evidence for this internalized grammar. Base-resolution gradient attribution co-localizes with empirical master TF binding sites, correlates globally with vertebrate evolutionary conservation, and dissects the modular assembly of shared regulatory domains such as *SLC25A25* and *ZMYND8*, where the model dynamically shifts sequence attention to distinct, lineage-defining TF motifs across cell types (**Fig. 3e**). Crucially, *in silico* constituent ablation establishes that Borzoi-SE captures higher-order constituent cooperativity and functional hierarchies rather than additive motif weights: across the human *α*- and *β*-globin LCRs, computational element masking independently prioritized HS2 as the dominant functional constituent whose ablation triggers a locus-wide collapse of predicted super-enhancer activity (**Fig. 4**). Our experimental validation via CRISPR interference (CRISPRi), ATAC-seq and RT-qPCR in living cells corroborated this regulatory hierarchy: silencing of HS2 precipitated a domain-wide collapse of chromatin accessibility and globin transcription, whereas targeting secondary constituents induced strictly focal reductions (**Fig. 4e-g**). These convergent findings confirm that Borzoi-SE learns non-additive regulatory cooperativity directly from primary sequence.

This ability to model non-linear sequence perturbations enables Borzoi-SE to transition from descriptive genomic mapping to prospective causal variant discovery. While traditional pipelines identify LoF variants by overlapping mutations with pre-existing open chromatin, identifying GoF mutations that nucleate *de novo* super-enhancers within quiescent chromatin remains a formidable challenge. Borzoi-SE addresses this gap *ab initio*: the model accurately recapitulates the lineage-restricted nucleation of a super-enhancer upstream of *TAL1* driven by a non-coding insertion in T-ALL (**Fig. 5**), and identifies cardiac variants that create *de novo* NR3C1 and GATA4 motifs to drive super-enhancer status transition (**Fig. 7 and Supplementary Fig. 5**). Furthermore, model-predicted perturbation magnitudes quantitatively calibrate empirical histone acetylation QTL (haQTL) effect sizes, effectively separating candidate functional mutations from co-inherited linkage disequilibrium hitchhikers. Orthogonal experimental evidence, spanning allele-specific chromatin accessibility, single-nucleus transcriptomics, dexamethasone-dependent glucocorticoid receptor recruitment, and iPSC-derived cardiomyocyte models, demonstrates that single-base alterations prioritized by Borzoi-SE directly initiate molecular cascades culminating in altered physiology (**Fig. 7**).

At the sequence level, base-resolution attribution elucidates the biophysical mechanism driving this activation, revealing a general non-linear regulatory phenomenon. Beyond imparting high positive attribution to the *de novo* MYB motif at *TAL1*, the micro-insertion dramatically elevated the attribution of an adjacent, endogenous PU.1/ETS motif that remained dormant in the wild-type sequence (**Fig. 5c**). Similarly, at the *CHRM2* locus, the *de novo* creation of an AR/NR3C1 motif concomitantly upregulated the sequence attribution of the neighboring, sequence-unaltered MEF2 motif (**Fig. 7b**). This observation indicates that Borzoi-SE captures authentic synergistic motif cooperativity rather than linear motif additivity. In unperturbed chromatin, isolated low-affinity or orphan motifs often remain functionally latent due to the energetic barrier of nucleosome occupancy and short TF residence times. The acquisition of an anchor motif serves as a catalytic nucleation seed, cooperatively licensing neighboring transcription factors to form stable multimeric assemblies and trigger a super-enhancer status transition. This finding mirrors recent multiscale TF footprint discovery^49^, which demonstrated that transcription factor binding footprints and local chromatin accessibility are non-locally determined by cooperative dependencies on adjacent sequence motifs^50^. The ability of Borzoi-SE to autonomously uncover how a focal non-coding alteration “awakens” dormant surrounding regulatory grammar underscores its capacity to model higher-order *cis*-regulatory syntax directly from primary DNA sequence.

Several biological and methodological frontiers define key priorities for future development. At the empirical level, because continuous regulatory strength is calibrated against ROSE-derived annotations, borderline elements near the geometric inflection point exhibit residual uncertainty, while evolutionary syntax divergence suggests that multi-species joint training will be valuable for cross-species generalization. More fundamentally, however, relying strictly on one-dimensional sequence leaves open the question of whether the model has truly internalized physical 3D genome architecture, such as loop extrusion and chromatin topological domains, or is predominantly exploiting local 1D epigenetic correlations. To address this, ongoing efforts are expanding Borzoi-SE into a unified multi-task framework that jointly predicts 1D functional readouts (e.g., PRO-cap, ChIP-seq) alongside higher-order 3D/4D spatial contacts (e.g., Hi-C, Micro-C, and HiChIP). Forcing shared representations to simultaneously satisfy both local regulatory activity and spatial proximity constraints will ensure the model captures the physical governing principles of genome folding. Ultimately, extending this biophysically grounded framework to single-cell multi-omics and diverse patient cohorts will broaden our capacity to decode non-coding disease variants and accelerate sequence-directed therapeutics.

## Methods

### Model architecture

Borzoi-SE formulates super-enhancer (SE) discovery as a multi-task genomic object detection task directly from primary sequence. Given an unsegmented 524,288-bp one-hot encoded DNA sequence, the network outputs continuous prediction tracks across a central 409,600-bp window divided into 1,600 bins of 256 bp for *C* = 90 human cell types and tissues simultaneously. The architecture comprises four sequential modules (Fig. 1a).

### Pretrained Backbone and Down-Sampler (Module 1)

Sequence embeddings are extracted using the pretrained Borzoi foundation model. For a 524,288-bp input, the Borzoi convolutional and transformer trunk produces 16,352 spatial positions at a 32-bp resolution across 1,536 latent channels (1,536 × 16,352). To match the broader domain-level scale of enhancer clusters, a depthwise-separable down-sampler—comprising a grouped 1D convolution (kernel 9, stride 8, 1,536 groups), batch normalization, and GELU activation—compresses these representations to 2,044 spatial positions at a 256-bp resolution (1,536 × 2,044).

### Parallel Encoding and Dilated Cluster head (Module 2)

The down-sampled representations branch into two parallel streams (Fig. 1a): **(1) Direct Context Stream**: Preserves the 1,536-channel sequence embeddings to retain local transcription factor (TF) motif features. **(2) Dilated Cluster head**: Models broad enhancer clustering. A 1 × 1 convolution projects the 1,536 channels to 512 channels (batch normalization, GELU, dropout 0.5), followed by six residual dilated convolution blocks with exponentially increasing dilations (*d* ∈ {1, 2, 4, 8, 16, 32} ; each block consists of two convolutions with a kernel size of three, batch normalization, GELU, and dropout 0.2). A final 1 × 1 convolution outputs 90-channel cluster logits (90 × 2,044). The dilation schedule expands the receptive field of the Cluster head to approximately 64.7 kb (253 bins), safely accommodating the classical 12.5 kb stitching window of the ROSE algorithm to model cluster nucleation while filtering background noise.

### Channel-Attentive Feature Fusion (Module 3)

To condition SE calling directly on cluster nucleation, the 1,536-channel context features and 90-channel cluster logits are concatenated along the channel dimension (1,626 × 2,044) and recalibrated via a Squeeze-and-Excitation block. Spatial global average pooling summarizes each channel, followed by a two-layer bottleneck MLP (reduction ratio 16, compressing 1,626 channels to 101 channels with GELU, then restoring 1,626 channels) and a sigmoid gating function. Element-wise multiplication of the resulting channel weights with the input tensor allows the network to dynamically balance localized TF motif syntax against broader cluster context.

### Super-Enhancer Head and Output Generation (Module 4)

The recalibrated representation is processed by an SE head to discriminate true super-enhancers from typical enhancer clusters. The SE head projects the 1,626 channels to 512 channels via a convolution with a kernel size of three, refines the features through four dilated residual blocks (*d* ∈ {1, 2, 4, 8}; local receptive field ∼ 16.1 kb / 63 bins), reduces them to 256 channels via a 1 × 1 convolution, and maps to the 90 output channels. The SE head is deliberately shallower than the Cluster head because SE calling acts primarily as a re-scoring of regional cluster predictions rather than independent feature extraction. Finally, the 2,044 positions are center-cropped to 1,600 bins (409,600 bp) to eliminate boundary padding artifacts. A sigmoid activation yields the final prediction tensor *̂Y* ∈ [0, 1]^90×1,600^, providing cell-type-specific SE probabilities aligned across the genome.

### Training data

#### ENCODE H3K27ac datasets

We assembled 90 human H3K27ac ChIP–seq datasets from the ENCODE portal (**Supplementary Table 1**). For each dataset we retrieved the ENCODE alignments of all available biological replicates together with the matched control alignment and merged the replicate alignments per dataset before peak calling with Sambamba (v1.0.1)^17,51^.

#### H3K27ac ChIP-seq peak calling

Merged alignments were restricted to the primary chromosomes of the assembly and reads overlapping blacklisted regions (ENCODE hg38 blacklist, 910 regions) were removed. H3K27ac peaks were then called with MACS2 (v2.2.9.1, --nomodel --extsize 150 --keep-dup all -q 0.01), using the matched control as background^52^. Across the 90 datasets, we obtained a median of 79,140 peaks per dataset (range 29,500–311,596) and a median fraction of reads in peaks (FRIP) of 0.370 (range 0.121–0.741).

#### Super-enhancer identification

Super-enhancers (SEs) were identified from the H3K27ac signal with ROSE (v1.3.2, https://github.com/stjude/ROSE), using the MACS2 peak calls as input, the H3K27ac and matched control alignments for signal quantification, and a stitching distance of 12.5 kb. Briefly, ROSE stitches nearby constituent peaks into stitched regions, ranks the stitched regions by control-subtracted H3K27ac signal, and calls SEs as the regions above the inflection point of the resulting signal-versus-rank curve; the threshold is therefore estimated per dataset rather than fixed across datasets.

#### Annotations used as training labels

Every stitched region of every dataset was carried forward as an annotation and classified as either an SE or a non-SE stitched region by the ROSE ‘isSuper’ flag. Annotations were stored per dataset and chromosome as genomic intervals, so that a training example is defined by sequence alone and the label is retrieved by interval lookup at training time.

For model training, interval annotations were rasterized into 256-bp bins and assigned soft labels: bins fully inside an SE were labelled 2, bins overlapping an SE boundary 1.75, bins inside a non-SE stitched region 1, bins overlapping the boundary of a non-SE stitched region 0.75, and all remaining bins 0. Because SE intervals are nested inside the stitched-enhancers, SE labels take precedence wherever the two overlap, so the two annotation classes are mutually exclusive in the final label matrix. Training examples were sampled from anchor intervals rather than from genes or fixed genomic tiling: stitched regions were merged when separated by less than 12.5 kb, and merged blocks longer than the model input were split into equal sub-blocks. In each epoch we sampled up to 200 SE anchors per dataset and an equal number of non-SE stitched-region anchors (1:1), plus half as many background anchors drawn uniformly from blacklist-free positions; anchors were jittered (±150 kb for annotation-derived anchors, ±50 kb for background anchors), and every window was required to fall entirely within a chromosome. Input sequences of 524,288 bp were one-hot encoded, and the model predicted 1,600 bins of 256 bp (409,600 bp) per dataset after center cropping; sequence and labels were reverse-complemented together with probability 0.5 during training.

### Chromosome splits

Model development used the 22 human autosomes only (sex chromosomes and unplaced scaffolds were excluded). Chromosomes were assigned to training, validation and test sets: chr9 and chr14 were used for validation, chr3 and chr11 were reserved for testing, and the remaining 18 autosomes were used for training (Split 0). All downstream performance analyses used held-out test chromosomes.

### Benchmark

#### Cross-chromosome partitioning and data independence

To rigorously assess model generalizability and prevent data leakage, we implemented a five-fold cross-chromosome evaluation framework (Splits 0–4). Benchmarks relying on a single fixed set of test chromosomes suffer from compositional biases stemming from chromosome-specific variations in gene density and GC content. To eliminate these artifacts, sex chromosomes (chrX and chrY) were excluded to avoid confounding effects from dosage compensation and sex-biased epigenetic regulation. In each fold, the 22 human autosomes were partitioned into mutually exclusive subsets: two autosomes designated for validation, two distinct held-out autosomes reserved for blind testing, and the remaining 18 autosomes allocated for model training. The five-fold chromosome allocation was structured as follows:

**Split 0**: Validation on chr9 and chr14; testing on chr3 and chr11.

**Split 1**: Validation on chr1 and chr15; testing on chr2 and chr12.

**Split 2**: Validation on chr4 and chr18; testing on chr5 and chr17.

**Split 3**: Validation on chr6 and chr19; testing on chr7 and chr20.

**Split 4**: Validation on chr8 and chr21; testing on chr10 and chr13.

All downstream benchmark models were independently trained, tuned, and tested across all five splits. Hyperparameter tuning and decision threshold optimization were strictly confined to validation chromosomes.

#### Baseline model implementations and retraining

We benchmarked Borzoi-SE against four published deep learning architectures for super-enhancer prediction: DeepSE, SENet, MuSE, and DcSE^6–9^. All baselines were standardized to their native 3-kb input scale and retrained using their published hyperparameter configurations and loss functions. All baseline models incorporated early stopping based on validation loss. In total, 20 pan-tissue union models (4 architectures × 5 splits) were trained for experiments 1 and 2, and 80 lineage-specific models (4 architectures × 4 cell types (K562, HepG2, WTC11, and GM12878) × 5 splits) were independently retrained for Experiment 3.

#### Unified multi-scale scoring metric

Evaluating 3-kb baseline classifiers alongside Borzoi-SE’s 524-kb receptive field across variable-length genomic domains requires strict scale alignment. Averaging model predictions across entire domains artificially penalizes localized models over non-peak intervals, whereas evaluating only central windows disregards constituent enhancer density. We therefore established a peak-restricted maximum scoring standard (prcmax) to compute domain-level scores uniformly across all models.

For any evaluated genomic domain *U* (stitched enhancer by ROSE), let *P_U_* denote the set of constituent narrowPeak summits identified by MACS2 within *U*. Domain-level scores were calculated through the following steps:

##### Constituent peak scoring

For baseline models, the prediction for peak *p* ∈ *P_U_* was computed over the 3-kb window centered at the peak summit:

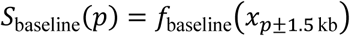

matching their native training resolution.

For Borzoi-SE, the score for peak *p* was calculated as the spatial average over all 256-bp bins *B_p_* spanned by the peak:

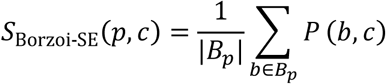

where *P*(*b*, *c*) is the predicted probability for bin *b* in cell channel *c*. For pan-tissue union tasks, scores were collapsed across channels via channel-wise maximum:

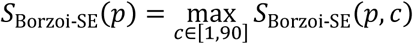

##### Domain-level aggregation

The final scalar score for domain *U* was assigned as the maximum score among its constituent peaks:

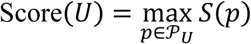

#### Experiment 1: Cluster-level pan-tissue binary classification

Experiment 1 benchmarked classification performance on discrete, pre-segmented ROSE enhancer clusters on held-out test chromosomes. Positive samples comprised all genomic intervals annotated as super-enhancers by the ROSE algorithm across any of the 90 reference cell types (isSuper = 1, merged with zero-gap distance). Negative samples comprised typical enhancer clusters (isSuper = 0) from which all positive SE regions were removed through base-pair-level subtraction. The test sets preserved the natural genomic imbalance of SEs to TEs (∼1:10) without artificial down sampling. Borzoi-SE extracted predictions from its native 524-kb receptive field centered at each cluster, whereas the 20 pretrained union baseline models scored the constituent peaks of each cluster using local 3-kb windows. All predictions were aggregated via the prcmax protocol.

Models were evaluated using the area under the receiver operating characteristic curve (ROC-AUC) and the area under the precision-recall curve (PR-AUC). In addition, validation-tuned F1 scores (F1_vt_) were determined by sweeping threshold *T* ∈ [0.01,0.99] on validation chromosomes:

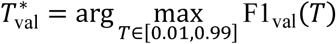

This threshold was subsequently frozen and applied directly to the held-out test set.

#### Experiment 2: Whole-chromosome end-to-end object detection

Experiment 2 challenged models to perform *de novo* regulatory domain detection directly across continuous, unfragmented held-out test chromosomes without prior knowledge of candidate cluster boundaries.

Continuous prediction tracks were generated across unsegmented test chromosomes: **Borzoi-SE** was evaluated via a sliding window of 524,288 bp with a step size of 204,800 bp. Valid central predictions spanning 409,600 bp (1,600 bins of 256 bp) were stitched across steps by taking the maximum probability across overlapping windows for each bin. The pan-tissue union track was derived by taking the maximum across all 90 cell-type channels:

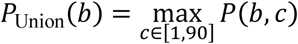

Baseline models were evaluated using a 3,000-bp sliding window stepped at 1,000-bp intervals (2,000-bp overlap). Window-level scalar predictions were projected onto the corresponding 256-bp bins, taking the maximum value across overlapping intervals to construct a continuous chromosome-wide prediction track.

To prevent sequencing artifacts and non-mappable assembly gaps from biasing evaluations, genomic regions overlapping ENCODE blacklists and contiguous stretches of assembly gaps (*N* bases in the GRCh38 primary assembly) were masked prior to metric calculation.

Detection fidelity was quantified across two distinct analytical tiers:

**Tier 1** (bin-level resolution): Evaluated continuous signal discrimination across all valid 256-bp bins outside masked regions using ROC-AUC and PR-AUC. Under pan-tissue union conditions, positive SE bins accounted for approximately 34%–39% of the unmasked genomic background. **Tier 2** (region-level object detection): Continuous prediction tracks were binarized across a threshold grid *T* ∈ [0.01,0.99]. Contiguous bins with *P*(*b*) ≥ *T* were merged, stitched across gaps ≤ 12.5 kb (50 bins), and filtered to remove spurious fragments < 3 kb, mirroring the empirical assembly criteria of the ROSE algorithm. Predicted domains *P_T_* were matched against ground-truth Union SE intervals GG using an intersection-over-union threshold IoU ≥ 0.3, where

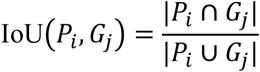

Candidate pairs with IoU ≥ 0.3 were sorted in descending order of IoU and matched greedily, removing both matched predictions and true regions to enforce a strict one-to-one mapping. Precision and recall were computed at each threshold *T* to generate region-level PR curves and calculate Region PR-AUC. The decision threshold *T*^∗^ maximizing region-level F1 on validation chromosomes was frozen and used to determine the final test Region F1 score. For all true-positive detections identified at an operating recall of 0.8, spatial boundary accuracy was quantified by computing the median IoU across matched prediction-ground truth pairs. Regulatory deserts were defined as unannotated genomic expanses lying outside the union of all 90 cell-type enhancer clusters, excluding ENCODE blacklists and assembly gaps:

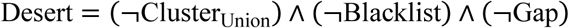

False-positive density was quantified at 80% recall as the total length (in megabases) of false-positive detections falling inside desert regions divided by the total megabases of desert sequence on the test chromosomes (Desert FP/Mb).

#### Experiment 3: Lineage-Specific Benchmark and Exclusivity Grammar

Experiment 3 investigated the capacity of each model to learn lineage-specific regulatory grammar across four representative human cell contexts spanning distinct embryonic lineages: K562 (erythroid), HepG2 (hepatocellular carcinoma), GM12878 (lymphoblastoid), and WTC11 (induced pluripotent stem cells). To ensure fair competition, all four baseline architectures were retrained on lineage-specific datasets across all five splits (4 models × 4 cell types × 5 splits = 80 retrained models).

**Positive training samples**: 3-kb windows centered on MACS2 narrowPeaks overlapping authentic SE intervals identified specifically within that cell type. **Negative training samples**: A 1:1 balanced mixture comprising non-SE peaks from the same cell type (MACS2 peaks located in typical enhancers) and random 3-kb unannotated genomic background regions lacking known peaks, blacklists, or assembly gaps, capped at 30,000 samples per training set.

Evaluations were performed across the full set of ROSE-annotated stitched enhancers identified within the target cell type (natural test prevalence: 4.0%–8.8% SEs). For baseline models, constituent peaks were scored using the corresponding lineage-retrained model and aggregated via prcmax. For Borzoi-SE, predictions were extracted directly from the dedicated output channel for that cell type, requiring no architectural fine-tuning or secondary retraining.

To determine whether models recognized lineage-defining sequence syntax rather than non-specific regulatory accessibility, we performed cross-lineage testing. Models trained on lineage *i* (or the dedicated channel *i* of Borzoi-SE) were evaluated against the held-out test cluster pools of lineage *jj*, generating a 4 × 4 cross-prediction matrix of test PR-AUC values. Diagonal entries (*i* = *jj*) quantified on-target lineage sensitivity, whereas off-diagonal entries (*i* ≠ *jj*) quantified cross-lineage false activation across non-target regulatory programs.

### Training

#### Two-stage training strategy

The adapter is trained in two stages, because the two output heads are randomly initialized while the Borzoi trunk is already informative: a first stage fits the task-specific heads with the pretrained trunk kept frozen and with a comparatively large learning rate, and a second stage then adapts the trunk itself with a much smaller learning rate, so that the pretrained sequence features are refined rather than overwritten. Both stages use the same data pipeline, the same anchor sampler, the same hierarchical loss and the same chromosome split, and both save the checkpoint with the best validation super-enhancer PR-AUC (**Supplementary Table 4**).

#### Stage 1 head-only training from random initialization

In the first stage the pretrained Borzoi backbone was loaded and frozen, so that the only trainable parameters were the randomly initialized adapter modules. Training therefore optimizes the mapping from frozen pretrained embeddings to the two annotation classes without perturbing the trunk, which is the only regime in which a large learning rate can be used safely: the adapter was optimized with AdamW at a head learning rate of 1 × 10^−4^ (weight decay 1 × 10^−2^) together with the hierarchical loss weight λ = 0.25 and a batch size of 12.

##### Stage 2: low-learning-rate fine-tuning of the trunk

The second stage initialized the model from the stage-1 best checkpoint and then unfroze the last transformer layer of the Borzoi backbone together with the decoder blocks that follow it. Keeping the large learning rate in this regime would destroy the pretrained representation, so the learning rate was reduced by two orders of magnitude: the adapter was optimized at 1 × 10^−6^ and the unfrozen backbone parameters at 2 × 10^−7^ with weight decay 1 × 10^−6^, while the loss weighting (λ = 0.25), batch size (12) and anchor sampling were unchanged. Stage 2 ran for 9 epochs and was terminated by early stopping; the best validation checkpoint was obtained in the first epoch, indicating that the trunk adaptation converged immediately from the stage-1 solution, and performance then remained essentially flat.

### Input sequence attribution

#### Attribution target

To explain model predictions, the target scalar was defined by summing SE-head logits of the channel of interest across the 256-bp bins covering the target enhancer. Taking the gradient of this region-level sum with respect to the one-hot input provides robustness against exact bin boundaries and directly matches the domain-level quantity predicted by the model.

#### Large-scale motif discovery by TF-MoDISco

Across all 90 channels, candidate windows with predicted SE probability > 0.5 overlapping annotated ROSE super-enhancers were selected. For each qualifying window, hypothetical contribution scores (gradients of the region-level score with respect to all four possible nucleotides across the 4 × 524,288 input) were computed and clustered using TF-MoDISco (v2.5.2; parameters: -n 20000 -w 100000)^53^. Discovered motif patterns were matched against annotated TF binding profiles using the built-in TOMTOM utility against the curated MotifCompendium human reference database^54,55^.

#### Specific sequences and variants (DeepLIFT)

For individual loci and genetic variants, DeepLIFT (via Captum) was used to avoid gradient saturation and yield signed, per-base attribution scores^30^. Attribution was calculated against 20 dinucleotide-shuffled sequence baselines, averaging the resulting multipliers before computing projected per-base scores. Reference and mutant alleles shared identical shuffled baselines to ensure that attribution differentials strictly reflected genetic variants rather than baseline stochasticity.

### Evolutionary conservation analysis

To evaluate whether Borzoi-SE-prioritized sequence motifs reflect functional constraint under purifying selection, base-resolution sequence attribution profiles were benchmarked against vertebrate evolutionary conservation. For each of the 90 modeled cell types, Gradient × Input attribution scores were computed across active super-enhancer domains. To mitigate single-nucleotide attribution variance, raw attribution magnitudes were smoothed using a 10-bp sliding window with a 5-bp step size. Base-by-base evolutionary conservation scores across 100 vertebrate species were extracted from the UCSC genome browser phastCons track^56^. For each cell type, genomic windows were stratified into deciles based on evolutionary conservation, and the concordance between mean binned attribution magnitude and phastCons scores was evaluated using Pearson correlation coefficients (r), with 95% confidence intervals estimated across all qualifying regulatory bins.

### Top-K candidate prioritization and ROSE rank fidelity

To evaluate prospective super-enhancer discovery, Borzoi-SE predictions were ranked based on maximum predicted probability. Precision@K and Core_Recall@K were computed across rank cutoffs K ∈ [1, 50]. Core super-enhancers were defined as the top 50% biologically dominant SEs ranked by empirical H3K27ac ChIP-seq signal intensity within each cell type. To determine whether predicted probabilities quantitatively reflect biological regulatory potency, predicted SE probabilities were summed across the spatial bins of each stitched enhancer cluster:

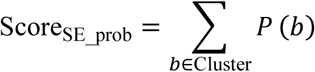

Spearman rank correlation coefficients (ρ) were calculated between this computational proxy and the empirical rank assigned by the ROSE algorithm across all 90 cell types.

### Cross-species zero-shot generalization in mouse

Cross-species sequence modeling was evaluated using matched primary tissues and developmental stages between human and mouse. H3K27ac ChIP-seq data and matched input controls for mouse cell lines, primary organs, and embryonic stages were obtained from the ENCODE project (**Supplementary Table 3**). Super-enhancers and typical enhancer clusters were called on the GRCm38 reference genome using MACS2 and ROSE under identical parameters (12.5-kb stitching distance). Full-length mouse genomic sequences (524,288 bp) centered on target loci were extracted and supplied directly to the human-trained Borzoi-SE deployment model without secondary parameter training or domain adaptation. Predicted outputs from corresponding human channels were evaluated against mouse experimental ROSE annotations to compute pairwise PR-AUC and ROC-AUC values.

### Cell culture and CRISPR interference

K562 cells were cultured in RPMI 1640 medium (GIBCO) and HEK293T cells were cultured in DMEM (GIBCO, C11995500BT) supplemented with 10% FBS (Noverse, NFBS-2500) and 1% (100 U/mL) Penicillin-Streptomycin (GIBCO, 15140122) at 37°C with 5% CO2. CRISPR interference was used to repress target enhancers. sgRNAs were designed against target regions and cloned into the pLentiGuide-sgRNA-Puro vector (Addgene, 52963). Correct plasmid insertion was verified by Sanger sequencing. The pLentiGuide-sgRNA-Puro plasmid and the pLV-dCas9-KRAB-MeCP2-EGFP plasmid were packaged into lentiviruses in HEK293T cells using lentiviral packaging system with psPAX2 and pMD2.G. Viral supernatants were collected at 48 and 72 h after transfection. K562 cells were co-infected with sgRNA and dCas9-KRAB-MeCP2 lentiviruses. For each enhancer, at least 2 sgRNAs were used individually to target the element. The gRNA sequences used in the study are: HS1 (TGAGGGAGGGGCCTAGCTAA; GGGACTGAGAAGGCAATAGC); HS2 (TAGAGTGATGACTCCTATCT; AACCTTCTAAGCAAACCTTC; TGCCCTGTAAGCATCCTGCT; GCCCTGTAAGCATCCTGCTG); HS3 (CTGCCAGCCTATAACCCATC; TGCCAGCCTATAACCCATCT); HS4 (AGTCCACCCCTTCTCGGCCC; AGTGAGTCAGCCTAGTAGAG); HS5 (GCTGCTGTTATGACCACTAG; AGTCCTGCCAGATATAGGTC). At 48 h after infection, EGFP-high cells were isolated by FACS. Cells were then selected with puromycin and subjected to downstream ATAC-seq and RT-qPCR analyses.

### ATAC-seq profiling and data processing

ATAC-seq was performed using 50,000 K562 cells using standard omni-ATAC method^57^. Cell pellets were washed with ice-cold PBS, resuspended in 50 μL of freshly prepared lysis buffer, and incubated on ice for 3 min. Lysis was stopped by adding 1 mL resuspension buffer, followed by gentle mixing. Nuclei were collected by centrifugation at 500 × g for 10 min at 4°C. The pellet was resuspended in 50 μL transposition reaction buffer containing assembled Tn5 transposase (Vazyme, S601) and incubated at 37°C with shaking at 1,000 rpm for 1 h. DNA was purified using a PCR purification kit (Qiagen, 28104) and amplified using HiFi HotStart Ready Mix DNA Polymerase (KAPA, KK2602) with indexed primers containing i5 and i7 adapters. PCR products were purified. Libraries were quantified using a Qubit fluorometer and subjected to paired-end sequencing.

ATAC-seq raw reads were trimmed and quality-filtered using fastp (v0.23.2), followed by paired-end alignment to the reference genome with Bowtie2 (v2.4.5) (--end-to-end --very-sensitive -X 2000, --no-mixed --no-discordant)^58,59^. Reads mapped to canonical chromosomes were retained, while non-canonical contigs were discarded with Sambamba (v1.0.1)^51^. PCR duplicates were marked and removed, and low-quality/blacklist-contaminated reads were further filtered by SAMtools (v1.15)^60^. Normalized chromatin accessibility tracks were generated via deepTools (v3.5.1) bamCoverage with RPGC normalization and 100-bp read extension^61^. Accessible peaks were identified using MACS2 (v2.2.7.1) in BAMPE mode (q-value < 0.01)^52^.

### *In silico* constituent ablation and locus activity integrals

To model higher-order hierarchies across compound regulatory domains, we performed systematic in silico sequence ablation across the human α-globin (chr16:60,000–190,000) and β-globin (chr11:5,100,000–5,350,000) locus control regions (LCRs). Individual constituent enhancers, promoters, and structural boundaries were computationally ablated by masking their native genomic sequences with ’N’ characters (encoded as all-zero one-hot vectors). The wild-type (WT) and in silico ablated mutant (MUT) sequences were evaluated by Borzoi-SE to compute per-bin super-enhancer probability deltas:

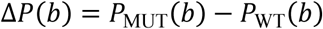

To quantify locus-wide regulatory consequences, we computed the locus activity integral ratio across the continuous domain:

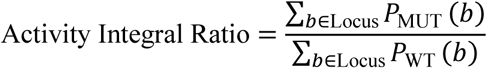

A ratio near 1.0 reflects functional dispensability, whereas substantial depletion (< 0.85) establishes the target element as an indispensable structural or functional hub.

### In silico variant screening and haQTL calibration

For genome-wide variant screening, genetic variants cataloged within ±10 kb of active enhancers across brain, heart, and muscle systems were evaluated. Sequences centered on the variant position (524,288 bp) containing either the reference or alternative allele were scored to compute the peak probability delta (Δ). Variants were categorized into: candidate Gain-of-Function (GoF; Δ ≥ 0.025), candidate Loss-of-Function (LoF; *P*_WT_ ≥ 0.30 and Δ ≤ −0.025), and neutral.

Computational Δ effect sizes were benchmarked against empirical histone acetylation quantitative trait loci (haQTLs) effect sizes (Beta) from corresponding tissues^36^.

### Motif scanning and transcription factor annotation

*De novo* motif patterns extracted by TF-MoDISco (v2.5.2) were matched against the curated human motif collection from the MotifCompendium package^19,55^. For targeted scanning of prioritized regulatory sites (e.g., *CHRM2* locus), position weight matrices (PWMs) were scanned across reference and mutant sequence windows using FIMO (*p* < 1e-4, v5.0.2) against the HOCOMOCO v14 database to determine the exact coordinates and motif integrity of disrupted or *de novo* created factor binding sites^54,62^.

### WASP allele-swap correction and allelic quantification

Allele-specific evidence was collected for the two Borzoi-SE gain-of-function candidates: chr7:136884669_T_C (rs1424569) and chr16:74903324_C_A (rs9937638). ENCODE heart (left/right ventricle) ATAC-seq datasets (all paired-end) were used: chr7: ENCSR745CGG, ENCSR286STX, ENCSR451JSB, ENCSR374ZRW, ENCSR439TZT, ENCSR053SGP, ENCSR327DCG; chr16: ENCSR846VPV, ENCSR769DGC, ENCSR133CMC, ENCSR491VKV. ENCODE heart 10X Genomics snRNA-seq datasets (single-end; read 1 carries the 16-bp cell barcode and the 12-bp UMI) were used: ENCSR349AHE, ENCSR079LVG, ENCSR630LZS, ENCSR851JBE, ENCSR151ONW, ENCSR763BII, ENCSR002SMQ, ENCSR753YOZ, ENCSR813PWQ. iPSC-derived cardiomyocyte ChIP-seq datasets (all single-end) from GEO GSE85631 were used: H3K27ac in four lines (2_6, 4_2, 5_6, MUT) for chr7 and GATA4 in four lines (1_8, 4_2, 5_6, WT) for chr16^45^. U2OS GR ChIP-seq datasets (all single-end) from GEO GSE274344 were used: Dex-treated, DMSO-treated and input^46^.

Heterozygosity at the variant position was established from the reads themselves, pooling all sequencing data of the same individual (for the iPSC-cardiomyocyte lines: H3K27ac, H3K27me3, H3K36me3, H3K4me3, Med1, Tbx5, GATA4 and input ChIP-seq datasets; for U2OS: Dex, DMSO and input ChIP-seq datasets; for each ENCODE experiment, technical-replicate libraries had already been merged). A site was called heterozygous when at least two reads carrying the reference allele and at least two reads carrying the alternative allele were present with base quality ≥ 20 and mapping quality ≥ 20. Note that heterozygosity was deliberately defined by the presence of both alleles rather than by an allele fraction close to 50%. Because the working hypothesis is that the alternative allele is functional (more accessible chromatin, more nascent transcription, more TF binding), the phenotype itself shifts the allele fraction; using the deviation from 50% to reject heterozygosity would therefore be circular.

For every sample, the observed allele fraction was corrected for reference-mapping bias with the WASP allele-swap procedure^47^. ATAC-seq BAMs were processed with the standard WASP2 pipeline (v1.4.1; https://github.com/mcvickerlab/WASP2). To eliminate reference mapping bias, allele-specific reads were filtered using the WASP2 pipeline (wasp2-map). For each sample, alignments across the target chromosome were extracted, and read pairs overlapping heterozygous sites were identified. Haplotype-swapped reads were generated using wasp2-map make-reads and realigned to the reference genome using Bowtie2 in end-to-end mode (-k 5 -X 2000 --mm). Finally, wasp2-map filter-remapped was applied to retain only read pairs that consistently returned to their exact original coordinates, which were then merged with reads requiring no remapping to generate the bias-corrected BAM file. WASP2 accepts paired-end input only. However, the ChIP-seq and 10x Genomics snRNA-seq alignments used here were treated as single-end, so these two assays were corrected with WASP implementations that operate on single-end reads. For snRNA-seq, the WASP implementation built into the STAR (v2.7.11a) aligner was used (--waspOutputMode SAMtag and --varVCFfile)^63^. Reads overlapping the variant were extracted from the ENCODE STARsolo alignments, realigned with STAR against the GRCh38 reference genome, and only alignments that passed the WASP test were retained. The single-end ChIP-seq libraries were corrected with the same single-end swap logic applied with the aligner that had produced the ChIP alignments (bowtie2 --end-to-end --very-sensitive)^58^. Reads overlapping the variant were rewritten into the other haplotype, realigned with identical settings, and retained only when the swapped copy mapped to the same coordinate; reads whose swapped copy moved or failed to align were discarded.

### Visualization and statistical analysis

Data visualization and statistical analyses were performed in Python using Matplotlib (v3.10.0) and Seaborn (v0.13.0) for graphical rendering, and SciPy (v1.15.0) for statistical testing.

### Use of Large Language Models (LLMs)

LLMs, such as ChatGPT, were used to assist with language editing and improve readability. The authors take full responsibility for the final manuscript.

## Data availability

The raw and processed ATAC-seq data generated in this study for the human β-globin LCR CRISPRi experiments in K562 cells have been deposited in the NCBI Gene Expression Omnibus (GEO) under accession number GSE347690. The human and mouse H3K27ac ChIP-seq datasets and matched control BAM files used for model training and benchmark evaluations were obtained from the ENCODE portal (**Supplementary Table 1 and 3**). Orthogonal validation ChIP-seq datasets in K562 cells for GATA1 (ENCSR000EFT), CTCF (ENCSR000AKB), NFE2 (ENCSR552YGL), along with baseline K562 ATAC-seq (ENCSR868FGK), were retrieved from ENCODE. Population genetics and functional QTLs: human tissue haQTL and eQTL summary statistics were downloaded from Zenodo (https://doi.org/10.5281/zenodo.7992724) and the GTEx Portal, respectively. Orthogonal multi-omics cardiac cohorts: primary human heart heterozygous donor ATAC-seq and single-nucleus RNA-seq (snRNA-seq) datasets were obtained from the ENCODE consortium (**Supplementary Table 5**). Dexamethasone-treated U2OS GR ChIP-seq datasets are available under GEO accession GSE274344. iPSC-derived cardiomyocyte ChIP-seq datasets (GATA4, H3K27ac, and H3K4me3) across varied genotypes are available under GEO accession GSE85631.

## Code availability

The complete source code and pretrained model for Borzoi-SE are publicly available on GitHub (https://github.com/Liangyan-1996/Borzoi-SE) and Zenodo (https://zenodo.org/records/22840452)^64^. The benchmark execution pipelines and reproduction scripts for DeepSE, SENet, MuSE, and DcSE are documented within the same repository.

## Acknowledgements

We thank Dr. Xushen Xiong and Dr. Ning Shen for guidance and discussions on model development. We thank Kailong Li for sharing dCas9-KRAB-MeCP2. We also acknowledge the support from the core facilities and computing platform of Liangzhu Laboratory at Zhejiang University School of Medicine.

## Fundings

The current work is supported by National Natural Science Foundation of China 82370123, 32350023 (to N.L.), Zhejiang Provincial Natural Science Foundation R25C060004 (to N.L.), Zhejiang Provincial Department of Science and Technology 2024SSYS0020 (Liangzhu laboratory, to N.L.), and Key R&D Program of Zhejiang (Research on Innovative Drug Development and Precision Clinical Translation for Genetic Mutation-Related Diseases).

## Author contributions

N.L. and Y.L. conceived the research. Y.L. developed the model and performed computational experiments. Y.L., X.X., Y.M. and Z.Z. analyzed data and generated figures. X.X. performed laboratory experiments. N.L., Y.L., X.X., Y.M. and Z.Z. wrote the manuscript. All authors reviewed and approved the final manuscript.

## Competing interests

The authors declare no competing interests.

---

**Supplementary Figure 1:** Extended cross-chromosome benchmark evaluations. **a**, Pan-tissue binary classification on pre-stitched enhancer clusters (Experiment 1): receiver operating characteristic (ROC) curves (left) and cluster-level AUROC bar plots (right). **b**, Median intersection-over-union (IoU) of true-positive SE detections at 80% recall on held-out test chromosomes (Experiment 2). **c**, Lineage-specific AUROC (left) and validation-tuned F1 scores (right) across four representative cell lines (Experiment 3). **d**, Pairwise cross-lineage evaluation matrices (4 × 4) showing mean AUPRC across five splits for Borzoi-SE and four baseline models across predicted (rows) and ground-truth (columns) lineages. For **a**–**c**, curves and bar heights denote the mean across *n* = 5 independent chromosome splits; shaded ribbons and error bars denote s.e.m.; individual points represent separate splits.

**Supplementary Figure 2:** Extended model performance metrics, rank fidelity, and cross-channel specificity across 90 human cell types. **a**, Super-enhancer detection precision versus recall across all *n* = 90 cell types on held-out test chromosomes (Split 0: chr3 and chr11). The dashed contour marks the F1 = 0.5 boundary; marker size and color intensity reflect the number of ground-truth SEs per cell type; the red star denotes global micro-average performance. **b**, Detection recall of Borzoi-SE (mean ± s.d.) stratified across deciles of biological SE strength (Decile 1 = top 10% highest H3K27ac intensity; Decile 10 = lowest 10%). The red curve indicates micro-average recall across deciles. **c**, Cumulative predicted SE probability sum (*y*-axis) versus biological ROSE rank (*x*-axis; Rank 1 = strongest SE, inverted scale) across six representative cell types, highlighting super-enhancers (red) and typical enhancers (grey). **d**, Pairwise cross-channel evaluation matrix (AUROC) assessing Borzoi-SE predictions (rows) against experimental ground truth (columns) across all 90 cell types.

**Supplementary Figure 3:** *De novo* discovery of lineage-defining TF motif syntax and constituent enhancer dissection at the ZMYND8 locus. **a**, Top eight positive-contribution motif patterns discovered *de novo* by TF-MoDISco across five representative lineages: WTC11 (induced pluripotent stem cells), H9 (embryonic stem cells), K562 (erythroid), CD14⁺ monocytes (myeloid), and primary hepatocytes (hepatic). **b**, Multi-lineage genomic snapshot across the *ZMYND8* super-enhancer locus (chr20:47,313,226– 47,373,327; ∼60.1 kb), displaying H3K27ac ChIP-seq coverage and ROSE SE annotations across six cellular contexts (WTC11, GM23338, K562, CD14⁺ monocytes, neutrophils, and hepatocytes). Constituent enhancers 20-1 and 20-2 are highlighted. **c**, Base-resolution DeepLIFT sequence attribution profiles across all six lineages at constituent enhancers 20-1 and 20-2.

**Supplementary Figure 4:** Borzoi-SE guides de novo identification and base-resolution dissection of functional loss-of-function regulatory variants. **a**, Multi-lineage genomic snapshot across the human *LDHA* locus (chr11:18,289,328–18,426,497). Top tracks depict predicted super-enhancer probability changes (Δ) across 12 human cell lines and primary tissues following the G>C transversion (chr11:18,369,690_G_C); bottom tracks display normalized baseline H3K27ac ChIP-seq coverage in wild-type cells. **b**, Multi-lineage genomic snapshot across the human *MAP2K6* locus (chr17:69,275,528–69,685,128). Top tracks depict predicted probability changes (Δ) across seven representative cell types following the G>A transition (chr17:69,480,329_G_A); bottom tracks display wild-type H3K27ac ChIP-seq coverage. **c**, Base-resolution DeepLIFT sequence attribution profiles across chr11:18,369,635–18,369,705 in gastrocnemius medialis, highlighting the disruption of the STAT3 motif and the adjacent master regulator MEF2C motif. **d**, Base-resolution DeepLIFT sequence attribution profiles across chr17:69,480,305–69,480,380 in gastrocnemius medialis, highlighting the disruption of the composite AR/NR3C1 and NFI motifs.

**Supplementary Figure 5:** Orthogonal validation of the chr16_74903324_C_A GoF variant. **a**, Multi-lineage genomic snapshot across the human *WDR59* locus (chr16:74,698,523–75,108,123). Top tracks depict predicted super-enhancer probability changes (Δ) following the C>A transversion (chr16:74,903,324_C_A) across 12 human cell types; bottom tracks display normalized baseline H3K27ac ChIP-seq coverage in wild-type cells. **b**, Base-resolution DeepLIFT sequence attribution profiles across chr16:74,903,270–74,903,370 in right atrial auricular tissue, highlighting the *de novo* creation of a GATA4 motif adjacent to an existing MEIS1/2 motif. **c**, Normalized H3K4me3, H3K27ac, and GATA4 ChIP-seq coverage in iPSC-derived cardiomyocytes across seven to nine cell lines (chr16:74,820,000–74,970,000) comparing heterozygous (orange) and homozygous wild-type (blue) lines. **d**, Allele-specific read pileup (± 15 bp around chr16:74,903,324_C_A) in heterozygous primary myocardium from ENCODE (left, ATAC-seq) and iPSC-derived cardiomyocytes (right, GATA4 ChIP-seq). **e**, Genome-wide ROSE ranking curves across individual iPSC-CM lines for chr7:136,884,669_T_C (top) and chr16:74,903,324_C_A (bottom). Stitched enhancer rank (*x*-axis) is plotted against total H3K27ac signal (*y*-axis), highlighting typical enhancers (grey) and *bona fide* super-enhancers crossing the inflection point (red).

## Notes

https://github.com/Liangyan-1996/Borzoi-SE

https://zenodo.org/records/22840452

### Competing Interest Statement

The authors have declared no competing interest.

## References

1. Whyte, W. A. et al. Master Transcription Factors and Mediator Establish Super-Enhancers at Key Cell Identity Genes. Cell 153, 307–319 (2013).

2. Bu, H., Hao, J., Gan, Y., Zhou, S. & Guan, J. DEEPSEN: a convolutional neural network based method for super-enhancer prediction. BMC Bioinformatics 20, 598 (2019).

3. Khan, A. & Zhang, X. Integrative modeling reveals key chromatin and sequence signatures predicting super-enhancers. Sci Rep 9, 2877 (2019).

4. Boeva, V. et al. Heterogeneity of neuroblastoma cell identity defined by transcriptional circuitries. Nat Genet 49, 1408–1413 (2017).

5. Lv, J. et al. 3D-super-enhancers are condensate-associated cis-regulatory communities. Nucleic Acids Res 54, gkag191 (2026).

6. Ji, Q.-Y., Gong, X.-J., Li, H.-M. & Du, P.-F. DeepSE: Detecting super-enhancers among typical enhancers using only sequence feature embeddings. Genomics 113, 4052–4060 (2021).

7. Zhang, A. & Jia, J. DcSE: an improved densenet with enhanced attention fusion for super-enhancer prediction. BMC Genomics 27, 428 (2026).

8. He, W., Zhou, H., Zuo, Y., Bai, Y. & Guo, F. MuSE: A deep learning model based on multi-feature fusion for super-enhancer prediction. Comput Biol Chem 113, 108282 (2024).

9. Luo, H. et al. SENet: A deep learning framework for discriminating super- and typical enhancers by sequence information. Comput Biol Chem 105, 107905 (2023).

10. Khan, A. & Zhang, X. dbSUPER: a database of super-enhancers in mouse and human genome. Nucleic Acids Res 44, D164–D171 (2016).

11. Kelley, D. R. et al. Sequential regulatory activity prediction across chromosomes with convolutional neural networks. Genome Res 28, 739–750 (2018).

12. Avsec, Ž., et al. Effective gene expression prediction from sequence by integrating long-range interactions. Nat Methods 18, 1196–1203 (2021).

13. Linder, J., Srivastava, D., Yuan, H., Agarwal, V. & Kelley, D. R. Predicting RNA-seq coverage from DNA sequence as a unifying model of gene regulation. Nat Genet 57, 949–961 (2025).

14. Avsec, Ž., et al. Advancing regulatory variant effect prediction with AlphaGenome. Nature 649, 1206–1218 (2026).

15. Jaganathan, K. et al. Predicting expression-altering promoter mutations with deep learning. Science eads7373 (2025) doi:10.1126/science.ads7373.

16. Mansour, M. R. et al. An oncogenic super-enhancer formed through somatic mutation of a noncoding intergenic element. Science 346, 1373–1377 (2014).

17. Dunham, I. et al. An integrated encyclopedia of DNA elements in the human genome. Nature 489, 57–74 (2012).

18. Popescu, D.-M. et al. Decoding human fetal liver haematopoiesis. Nature 574, 365–371 (2019).

19. kundajelab/tfmodisco. Kundaje Lab (2026).

20. Blayney, J. W. et al. Super-enhancers include classical enhancers and facilitators to fully activate gene expression. Cell 186, 5826–5839.e18 (2023).

21. Hay, D. et al. Genetic dissection of the α-globin super-enhancer in vivo. Nat Genet 48, 895–903 (2016).

22. Huang, J. et al. Dissecting super-enhancer hierarchy based on chromatin interactions. Nat Commun 9, 943 (2018).

23. Shin, H. Y. et al. Hierarchy within the mammary STAT5-driven Wap super-enhancer. Nat Genet 48, 904–911 (2016).

24. Hughes, J. R. et al. Annotation of cis-regulatory elements by identification, subclassification, and functional assessment of multispecies conserved sequences. Proceedings of the National Academy of Sciences 102, 9830–9835 (2005).

25. Li, Q., Zhang, M., Han, H., Rohde, A. & Stamatoyannopoulos, G. Evidence that DNase I hypersensitive site 5 of the human beta-globin locus control region functions as a chromosomal insulator in transgenic mice. Nucleic Acids Res 30, 2484–2491 (2002).

26. Wai, A. W. K. et al. HS5 of the human β-globin locus control region: a developmental stage-specific border in erythroid cells. EMBO J 22, 4489–4500 (2003).

27. Talbot, D. & Grosveld, F. The 5’HS2 of the globin locus control region enhances transcription through the interaction of a multimeric complex binding at two functionally distinct NF-E2 binding sites. EMBO J 10, 1391–1398 (1991).

28. Zhu, X. et al. A facilitated tracking and transcription mechanism of long-range enhancer function. Nucleic Acids Res 35, 5532–5544 (2007).

29. Cho, Y. et al. The role of transcriptional activator GATA-1 at human beta-globin HS2. Nucleic Acids Res 36, 4521–4528 (2008).

30. Shrikumar, A., Greenside, P. & Kundaje, A. Learning Important Features Through Propagating Activation Differences. Preprint at 10.48550/arXiv.1704.02685 (2019).

31. Bozhilov, Y. K. et al. A gain-of-function single nucleotide variant creates a new promoter which acts as an orientation-dependent enhancer-blocker. Nat Commun 12, 3806 (2021).

32. Christensen, A. H. et al. Gain-of-function enhancer variant near KCNB1 causes familial ST-depression syndrome. Eur Heart J 46, 3486–3497 (2025).

33. Prabhakar, S. et al. Human-Specific Gain of Function in a Developmental Enhancer. Science 321, 1346–1350 (2008).

34. Morris, J. A. et al. Discovery of target genes and pathways at GWAS loci by pooled single-cell CRISPR screens. Science 380, eadh7699 (2023).

35. Wu, Y. et al. Genome-wide fine-mapping improves identification of causal variants. Nat Genet 58, 940–951 (2026).

36. Hou, L. et al. Multitissue H3K27ac profiling of GTEx samples links epigenomic variation to disease. Nat Genet 55, 1665–1676 (2023).

37. GTEx Consortium. The GTEx Consortium atlas of genetic regulatory effects across human tissues. Science 369, 1318–1330 (2020).

38. Buniello, A. et al. Open Targets Platform: facilitating therapeutic hypotheses building in drug discovery. Nucleic Acids Res 53, D1467–D1475 (2025).

39. Zoodsma, M. et al. A genetic map of human metabolism across the allele frequency spectrum. Nat Genet 57, 2445–2455 (2025).

40. Graham, S. E. et al. The power of genetic diversity in genome-wide association studies of lipids. Nature 600, 675–679 (2021).

41. Christakoudi, S., Evangelou, E., Riboli, E. & Tsilidis, K. K. GWAS of allometric body-shape indices in UK Biobank identifies loci suggesting associations with morphogenesis, organogenesis, adrenal cell renewal and cancer. Sci Rep 11, 10688 (2021).

42. Ntalla, I. et al. Multi-ancestry GWAS of the electrocardiographic PR interval identifies 202 loci underlying cardiac conduction. Nat Commun 11, 2542 (2020).

43. Karczewski, K. J. et al. Pan-UK Biobank genome-wide association analyses enhance discovery and resolution of ancestry-enriched effects. Nat Genet 57, 2408–2417 (2025).

44. Hennis, K., Piantoni, C., Biel, M., Fenske, S. & Wahl-Schott, C. Pacemaker Channels and the Chronotropic Response in Health and Disease. Circulation Research 134, 1348–1378 (2024).

45. Ang, Y.-S. et al. Disease Model of GATA4 Mutation Reveals Transcription Factor Cooperativity in Human Cardiogenesis. Cell 167, 1734–1749.e22 (2016).

46. Sueyoshi, T. et al. Molecular interactions of glucocorticoid and mineralocorticoid receptors define novel transcription and biological functions. J Biol Chem 301, 108488 (2025).

47. van de Geijn, B., McVicker, G., Gilad, Y. & Pritchard, J. K. WASP: allele-specific software for robust molecular quantitative trait locus discovery. Nat Methods 12, 1061–1063 (2015).

48. Picard, D. & Yamamoto, K. R. Two signals mediate hormone-dependent nuclear localization of the glucocorticoid receptor. EMBO J 6, 3333–3340 (1987).

49. Hu, Y. Multiscale footprints reveal the organization of cis-regulatory elements.

50. Hu, Y. et al. Multiscale footprints reveal the organization of cis-regulatory elements. Nature 638, 779–786 (2025).

51. Tarasov, A., Vilella, A. J., Cuppen, E., Nijman, I. J. & Prins, P. Sambamba: fast processing of NGS alignment formats. Bioinformatics 31, 2032–2034 (2015).

52. Zhang, Y. et al. Model-based Analysis of ChIP-Seq (MACS). Genome Biol 9, R137 (2008).

53. Shrikumar, A., et al. Technical Note on Transcription Factor Motif Discovery from Importance Scores (TF-MoDISco) version 0.5.6.5. Preprint at 10.48550/arXiv.1811.00416 (2020).

54. Bailey, T. L., Johnson, J., Grant, C. E. & Noble, W. S. The MEME Suite. Nucleic Acids Res 43, W39–W49 (2015).

55. kundajelab/MotifCompendium. Kundaje Lab (2026).

56. Casper, J. et al. The UCSC Genome Browser database: 2026 update. Nucleic Acids Res 54, D1331–D1335 (2026).

57. Corces, M. R. et al. An improved ATAC-seq protocol reduces background and enables interrogation of frozen tissues. Nat Methods 14, 959–962 (2017).

58. Langmead, B. & Salzberg, S. L. Fast gapped-read alignment with Bowtie 2. Nat Methods 9, 357–359 (2012).

59. Chen, S. Ultrafast one-pass FASTQ data preprocessing, quality control, and deduplication using fastp. Imeta 2, e107 (2023).

60. Li, H. et al. The Sequence Alignment/Map format and SAMtools. Bioinformatics 25, 2078–2079 (2009).

61. Ramírez, F., Dündar, F., Diehl, S., Grüning, B. A. & Manke, T. deepTools: a flexible platform for exploring deep-sequencing data. Nucleic Acids Res 42, W187–W191 (2014).

62. Vorontsov, I. E. et al. HOCOMOCO in 2024: a rebuild of the curated collection of binding models for human and mouse transcription factors. Nucleic Acids Res 52, D154–D163 (2024).

63. Dobin, A. et al. STAR: ultrafast universal RNA-seq aligner. Bioinformatics 29, 15–21 (2013).

64. Liang, Y. Borzoi-SE: Clustered Enhancer Recognition via Borzoi-Embedded Representations of Underlying Sequences — code and model weights (v1.0.0). 10.5281/zenodo.22840452 (2026) doi:10.5281/zenodo.22840452.

