## Supplementary Figure 1-5 for "Decoding super-enhancer regulatory grammar and pathogenic gain-of-function variants directly from DNA sequence"

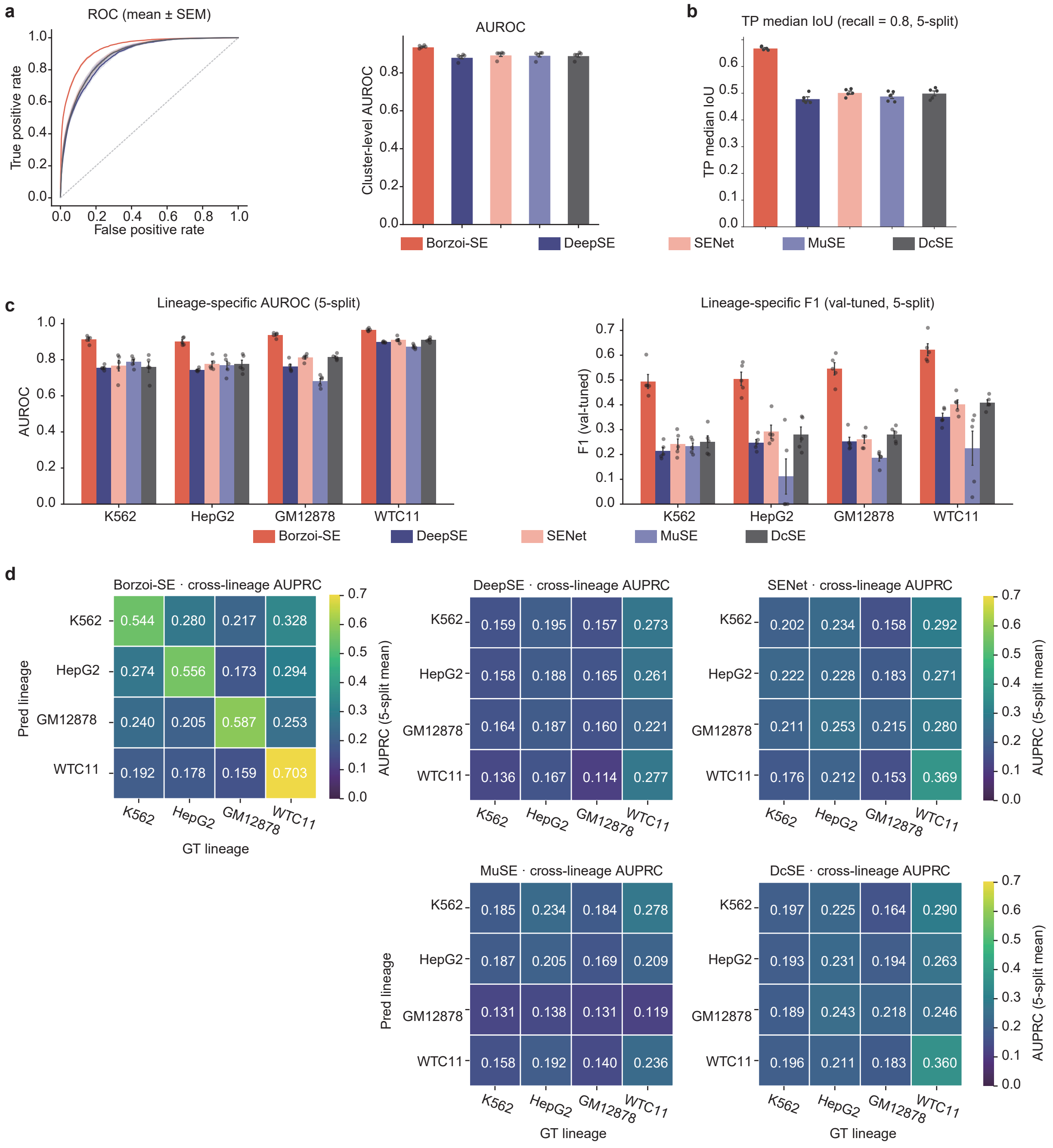

### Supplementary Figure 2

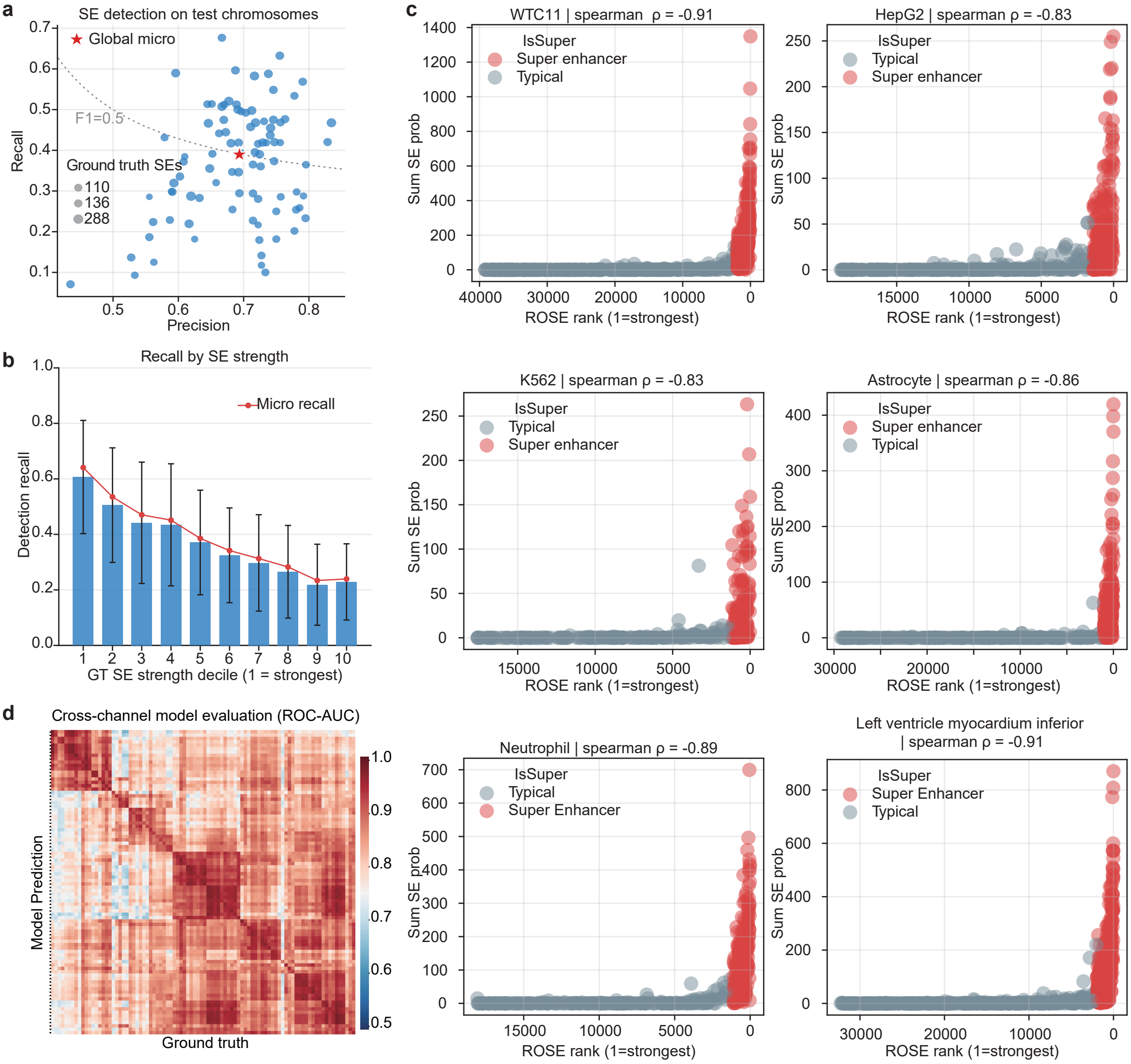

#### Supplementary Figure 3

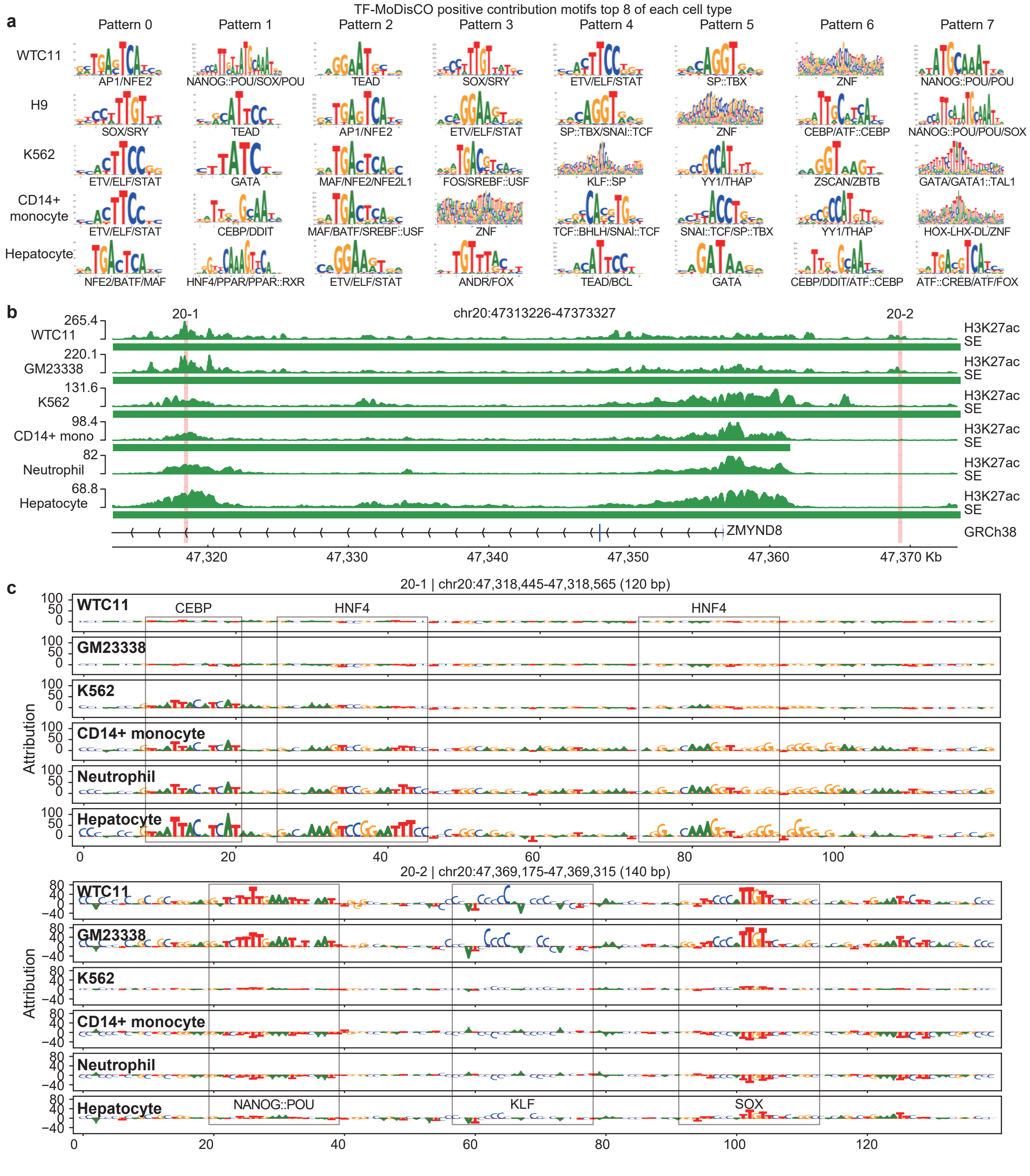

#### Supplementary Figure 4

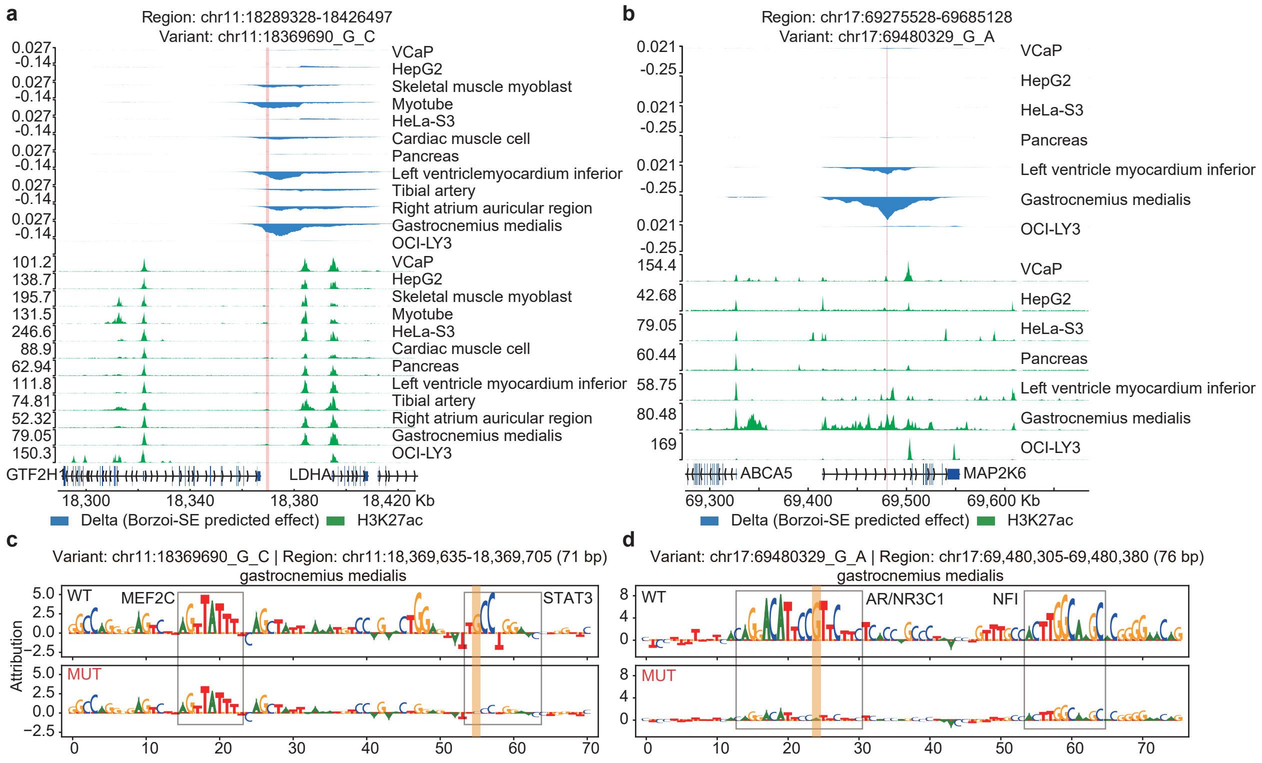

### Supplementary Figure 5

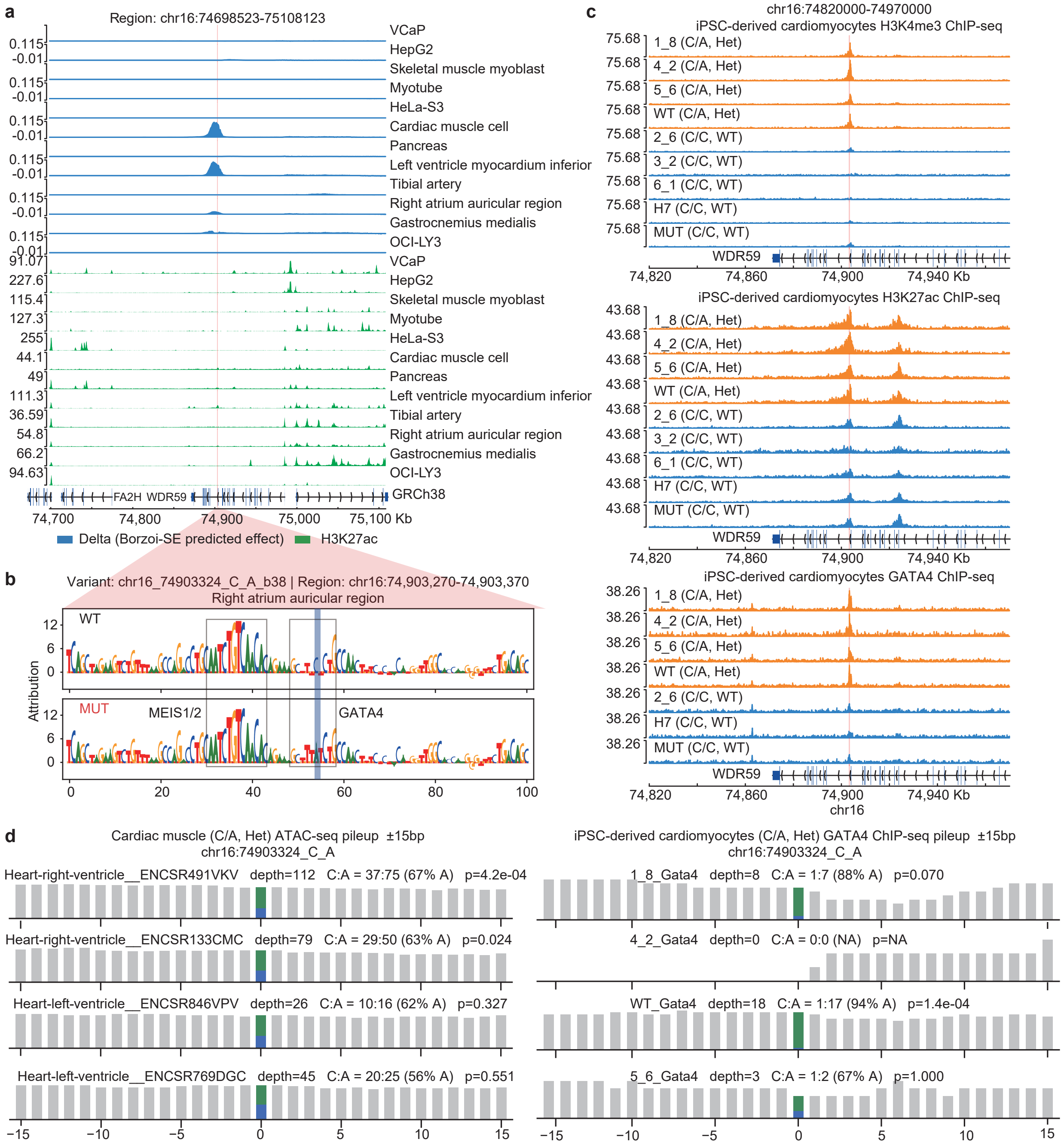

### Supplementary Figure 5 (continued)

**e**

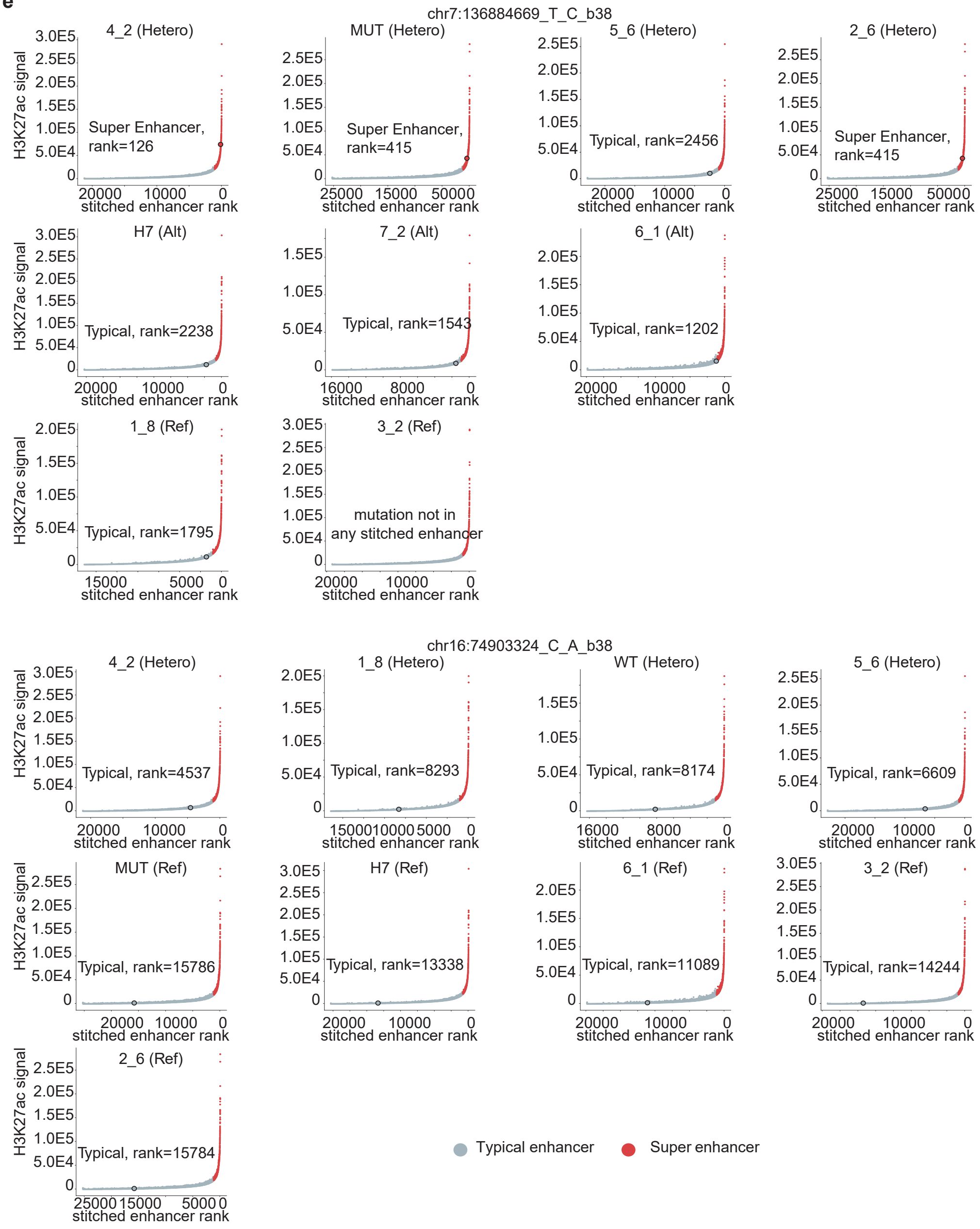
